# MAST1 is neuron-specific regulator of mitochondrial trafficking and abundance in human brain tissue

**DOI:** 10.64898/2026.11.04.764401

**Authors:** H.M. Arif Ullah, Kandy L. Napan, Ivan Khimach, Miguel A. Cuevas, Yongsoo Kim, Erin B. Taylor, Maria A. Serrano, Anthony S. Ricciardulli, Sophia Andreadis, Zhenni Jin, Nicole Losurdo, Andrew Harrahill, Colin Maguire, Jean J. Kim, Aleksandar Bajic, Wei Shen, Rong Sao, Pinar Bayrak-Toydemir, H. Joseph Yost, Yoshiho Ikeuchi, Jared Rutter, Lorenzo B. Botto, John C. Carey, Nichole Link, Alex Shcheglovitov

**Affiliations:** Neurobiology, University of Utah, Salt Lake City, UT 84112 USA; Biomedical Engineering, University of Utah, Salt Lake City, UT 84112 USA; Pathology, University of Utah, Salt Lake City, UT 84112 USA; Biochemistry, University of Utah, Salt Lake City, UT 84112 USA; Pediatrics, and University of Utah, Salt Lake City, UT 84112 USA; Psychiatry departments, University of Utah, Salt Lake City, UT 84112 USA; Cellular and Translational Research Core, Utah Clinical and Translational Science Institute, Salt Lake City, UT 84112 USA; Center for Regenerative Medicine, Department of Medicine, Boston University, Boston, MA; Advanced Technology Cores and Baylor College of Medicine, Houston, TX 77030, USA; Department of Molecular and Human Genetics, Baylor College of Medicine, Houston, TX 77030, USA; Jan and Dan Duncan Neurological Research Institute, Texas Children’s Hospital, Houston, TX 77030, USA; ARUP Laboratories, Salt Lake City, UT 84112 USA; Institute of Industrial Science, University of Tokyo, Tokyo 153-8505, Japan; Howard Hughes Medical Institute, University of Utah, Salt Lake City, UT 84112 USA

## Abstract

In human neurons, energy-demanding axonal and dendritic development requires sufficient mitochondrial distribution at distal locations, and the responsible mechanisms remain largely unknown. Here, we characterize rare *de novo* mutations in microtubule-associated serine/threonine kinase 1 (MAST1) in patients with multiple neurodevelopmental abnormalities, including microcephaly and thin corpus callosum. Experiments using induced pluripotent stem cell-derived telencephalic brain organoids and cortical neurons from patients and unaffected parents, as well as engineered lines demonstrate association of the *MAST1* mutations with organoid growth deficits and impaired neurites. Mechanistically, the *MAST1* mutations alter its interaction with mitochondria as well as mitochondrial trafficking and abundance, increasing the proportion of hyperactive mitochondria and reactive oxygen species accumulation. Exogenous wild-type MAST1 expression rescues neurite outgrowth in mutant neurons, as does prolonged treatment with creatine or *N*-acetyl-L-cysteine. This study identifies MAST1 as an important regulator of mitochondrial properties in human neurons and links microcephaly-associated *MAST1* mutations to neuronal mitochondrial dysfunction, impaired neurites, and neuronal loss.

## Introduction

Most aspects of neuronal development and survival are tightly regulated by energy availability^1–3^. Disrupted energy production and distribution are associated with neurological and psychiatric disorders^4,5^. Mitochondria play central roles in regulating neuronal energy production and distribution, including ATP delivery to distal locations in axons and dendrites to support energy-demanding processes such as membrane remodeling and neurite outgrowth^6^. From studies in model organisms, we know that anterograde and retrograde mitochondrial trafficking in neurites is mediated primarily by kinesin-1 and dynein–dynactin motors, respectively^7,8^. These motors are linked to mitochondria via adaptor proteins Miro1/2 and trafficking kinesin-binding proteins 1/2 (TRAK1/2) that also regulate mitochondrial targeting into axons and dendrites^8,9^. Upon delivery of mitochondria to specific locations in neurites, their localization is stabilized by the axonal Syntaphilins and dendritic vesicle-associated protein B, Rhotekin 2, and syndapin 1 protein complexes ^10–12^. Glucose, cytoskeletal architecture, and activity in the Liver kinase B1 (LKB1)–NUAK1 pathway are also known to regulate mitochondria motility in neurites^13–15^. Importantly, disruption of mitochondrial trafficking and stabilization impairs normal axonal or dendritic development^12,14,16^, and mutations in motor and adaptor proteins have been detected in patient with severe neurological disorders^17–20^. Given that human neurons have substantially longer and more complex neurites than neurons in model organisms, it is conceivable that additional proteins and pathways are involved in regulating mitochondrial trafficking and metabolism in human neurons^21,22^. Indeed, human-specific mitochondrial metabolism has been identified as the key regulator of the delayed morphological and functional maturation in human cortical stem cell-derived neurons^23^.

Microtubule-associated serine/threonine kinase 1 (MAST1) is expressed in neurons during brain development^24^. In addition to being enriched in the brain, it is also localized at both axons and dendrites where it interacts with microtubule-associated proteins to associate with microtubules^25^. MAST1 belongs to the AGC kinase family and contains an N-terminal domain of unknown function (DUF1908), a central AGC kinase domain, and a C-terminal PDZ domain^26^. Rare mutations predominantly in the domain of unknown function (DUF) and kinase domain of *MAST1* have been identified in patients with severe intellectual disability, autism, cerebral palsy, microcephaly, cortical malformations, and mega-corpus-callosum syndrome^24,27–34^. A mouse model carrying a patient-specific p.Leu278del mutation within the DUF domain exhibited an enlarged corpus callosum with increased numbers of midline-crossing axons, as well as elevated neuronal apoptosis, whereas *Mast1*-null mice lacked comparable morphological abnormalities^24^. More recently, it was demonstrated that phosphorylated MAST1 forms a complex with 14-3-3η, that Tau is a candidate MAST1 substrate, and that Tau phosphorylation is reduced in the p.Leu278del mouse model of mega-corpus-callosum syndrome^35^. In other studies, MAST1 and its homologs were found to regulate life span in *Caenorhabditis elegans*^36^, early cellularization in *Drosophila* embryo^37^, and cisplatin resistance in cancer cells^38^. Interestingly, the *Drosophila* homolog of MAST kinases, *dropout (dop)*, genetically interacts with components of the dynein–dynactin complex and promotes dynein-dependent apical transport of lipid droplets during cellularization ^37^, suggesting potential involvement of MAST kinases in regulating intracellular transport. In addition, an affinity-purification mass spectrometry screen using an exogenous, N-terminally tagged fragment of MAST1 in HEK293 cells demonstrated interaction of mitochondrial ribosomal proteins with MAST1^39^, raising the possibility of MAST1 association with mitochondria.

In the present study, we identified three unrelated pediatric patients with microcephaly, thin corpus callosum, and severe intellectual disability that each carry a rare *de novo* heterozygous missense mutation in the C-terminal region of *MAST1*. We generated induced pluripotent stem cells (iPSCs), iPSC-derived neurons, and brain organoids from two patients and two unaffected parents, as well as CRISPR-engineered human embryonic stem cell lines carrying a patient-specific mutation. We found that *MAST1*-mutant organoids exhibited microcephaly-associated growth deficits accompanied by altered cell composition, reduced neuropil, increased neuronal apoptosis, and impaired neurite extension. To identify the cellular and molecular mechanisms contributing to these deficits, we investigated the expression and subcellular localization of MAST1 in developing human neurons and organoids and found that MAST1 associates with mitochondria and is required for mitochondrial stabilization in neurites. *MAST1*-mutant neurons exhibited reduced MAST1 protein expression, diminished mitochondrial abundance, and altered mitochondrial trafficking in neurites, as well as elevated production of reactive oxygen species (ROS), impaired neurite outgrowth, and increased apoptosis. Importantly, exogenous MAST1 rescued neurite outgrowth deficits in MAST1-mutant neurons, and prolonged treatment with creatine or *N*-acetyl-L-cysteine (NAC) inhibited ROS production, increased neuronal survival, and improved neurite outgrowth, suggesting that up-regulation of MAST1 and targeting of bioenergetic and redox vulnerabilities in MAST1-mutant neurons offer potentially promising therapeutic strategies. Collectively, this study identifies MAST1 as an important regulator of mitochondrial trafficking and stabilization in developing human neurons and connects mitochondrial deficits to defective neurite outgrowth, neuronal loss, and microcephaly-associated organoid growth deficits.

## Results

### Organoids with MAST1 mutations exhibit microcephaly-associated growth deficits, increased neuronal apoptosis, reduced neuropil, and altered neural cell composition

We identified three unrelated individuals with microcephaly, thinning of the corpus callosum, developmental delay, and intellectual disability (**Figure 1A**; **Table S1**). Each individual carried a different rare de novo heterozygous missense variant in *MAST1*, encoding an amino acid substitution within the C-terminal region of the MAST1 protein (**Figure 1B**; **Table S2**). Analyses of the GTEx^40^ and PsychENCODE^41^ datasets showed that *MAST1* expression is enriched in the human brain relative to other tissues and its expression peaks during mid-gestation, respectively (**Figure S1**). To investigate the role of MAST1 in brain development and the pathogenesis of microcephaly, we initially used *Drosophila melanogaster* as a reliable in vivo model for brain development and microcephaly studies^42^. We reduced the expression of *dop*, a *Drosophila* homolog of human *MAST1*, in neural stem cells and neurons using four independent RNAi constructs targeting distinct regions of *dop* (**Figure S2A**). We observed no effects of *dop* knockdown on the size of *Drosophila* brain (**Figures S2B–S2E**). Moreover, expression of either wild-type (WT) human MAST1 or MAST1 carrying the patient-specific p.Pro1177Ser variant failed to rescue the lethality caused by complete loss of *dop*^37^. These results indicate that neural *dop* knockdown does not recapitulate MAST1-associated microcephaly and that human MAST1 cannot functionally substitute for *dop* in the developing *Drosophila* embryo, limiting the utility of *Drosophila* for modeling the effects of patient *MAST1* variants on brain development and suggesting potential species-specific differences in protein function, expression, localization, or interaction partners.

**Figure 1.**
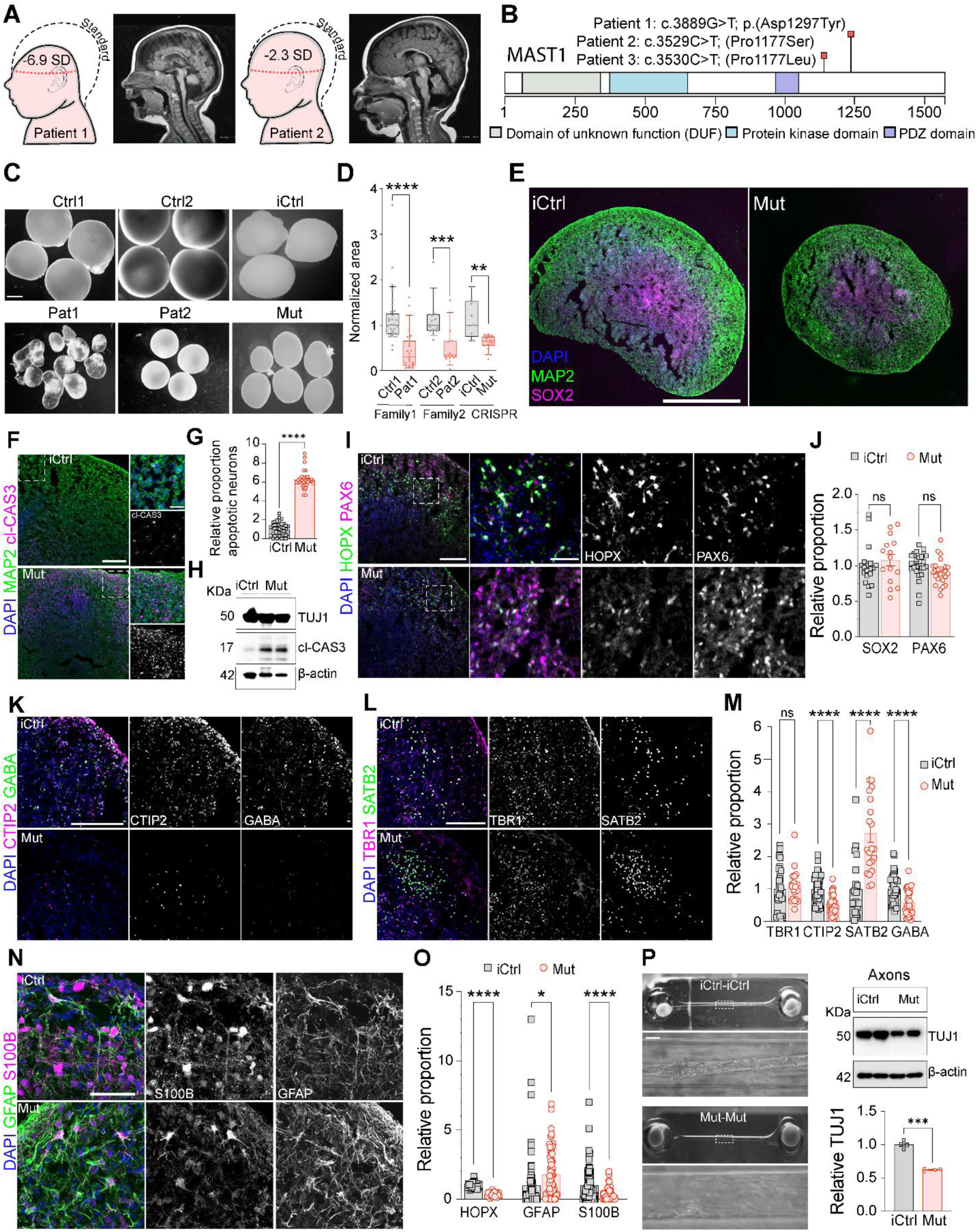
Organoid growth and cell composition deficits in 5-month-old organoids with MAST1 mutations. (A) Schematics of head circumference of each patient (left) and representative brain MRI images (right). (B) MAST1 protein domains and the *de novo* heterozygous MAST1 mutations identified in patients. (C) Images of 5-month-old SNR-derived brain organoids from control (ctrl1 and ctrl2), patient (pat1 and pat2), isogenic control (iCtrl), and MAST1 mutant (mut) lines. Scale bar: 1 mm. (D) Quantification of organoid size in 5-month-old ctrl1 (n=33), ctrl2 (n=17), pat1 (n=23), pat2 (n=13), iCtrl (n=9), mut1 (n=9) and mut2 (n=9) organoids produced in 3-7 batches. Dots: individual organoids. (E, F, I, K, L, and N) Images of 5-month-old iCtrl and MAST1 mut organoid sections immunostained for SOX2 and MAP2 (E), cleaved Caspase-3 and MAP2 (F), HOPX and PAX6 (I), CTIP2 and GABA (K), TBR1 and SATB2 (L), and GFAP and S100B (N) with DAPI. (G, J, M, and O) Normalized cell proportions in iCtrl and mutant organoids (n = 19 and 15 [SOX2], 40 and 35 [cl-Cas3/MAP2], 26 and 26 [PAX6], 27 and 20 [TBR1], 40 and 30 [CTIP2], 22 and 21 [SATB2], 40 and 30 [GABA], 40 and 30 [HOPX], 81 and 81 [GFAP], and 81 and 81 [S100β] randomly selected images obtained from 4 iCtrl and 3 mut organoids produced in 2 batches, respectively). (H) Immunoblots of TUJ1 and cl-Cas3 in 5-month-old organoid lysates obtained from iCtrl (n=3) and mut (n=6) organoids. β-actin was used as a loading control. (P) Images of pairs of iCtrl and mutant organoids in tissue chips with interconnected chambers and immunoblots with quantification of TUJ1 expression in lysates collected from connecting microfluidic channels (n = 4 channel lysates per group). β-actin was used as a loading control. Scale bars: 1 mm (E and P), 0.2 mm (F, I, K, and L), and 50 µm (N and zoom-in in F and I). Error bars = mean ± SEM. ns, not significant; *P<0.05, **P<0.01, ***P<0.001, and ****P<0.0001; unpaired two-tailed t-test.

To examine the effects of patient *MAST1* variants on human telencephalic brain development, we generated iPSCs from patients 1 and 2 as well as from two unaffected, sex-matched parents (**Figure S3**). In parallel, we used CRISPR-Cas9 editing to introduce the heterozygous p.Asp1297Tyr variant identified in patient 1, who demonstrated the most severe neurodevelopmental phenotype and microcephaly, into an independent control H9 embryonic stem cell line (**Figure S4**). We then generated single-neural-rosette (SNR)–derived telencephalic organoids according to our established protocol^43,44^ from matched control–mutant stem cell line pairs (**Figure S5**). At 4 months post-SNR isolation (5 months post-induction), SNR-derived organoids generated from both patient and engineered mutant lines were significantly smaller than matched control organoids (**Figures 1C and 1D**), indicating impaired organoid growth.

To assess the effects of MAST1 mutation on organoid cell composition, we examined cell type-specific marker protein expression in sections from H9 isogenic control (iCtrl) and engineered MAST1-mutant organoids by immunostaining (**Figures 1E–1O**). Immunostaining for the neuronal marker microtubule-associated protein 2 (MAP2) and neural progenitor (NP) marker SRY-box 2 (SOX2) revealed similar radial organization and proportions of neurons and progenitors in iCtrl and mutant organoids (**Figures 1E and 1F**). However, the absolute areas occupied by MAP2-and SOX2-expressing cells and number of cells were smaller in mutant organoid sections (**Figure 1E**). Mutant organoids contained neurons with fewer MAP2-expressing neurites (**Figure 1E**) and showed increased proportions of neurons expressing the apoptotic marker cleaved caspase-3 (**Figures 1F–1H**), suggesting that reduced neuropil and increased neuronal apoptosis may contribute to the impaired growth of MAST1-mutant organoids. Additionally, MAST1-mutant organoids were characterized by unchanged proportions of Paired Box 6 (PAX6)-expressing dorsal neural progenitors and T-box brain transcription factor 1 (TBR1)-expressing cortical neurons, and reduced proportions of homeodomain-only protein (HOPX)-expressing outer radial glia-like cells, COUP-transcription factor-interacting protein 2 (CTIP2)-expressing cortical neurons, gamma-aminobutyric acid (GABA)-expressing inhibitory neurons, and S100B-expressing astroglia (**Figures 1I–1O**). Interestingly, MAST1-mutant organoids showed elevated proportions of Special AT-rich sequence-binding protein 2 (SATB2)-expressing cortical neurons and glial fibrillary acidic protein (GFAP)-expressing astroglia (**Figures 1L–1O**), suggesting complex effects of MAST1 mutations on the specification and maintenance of neural cells in developing human telencephalic brain tissue.

While depleted proportions of HOPX-, CTIP2-, and GABA-expressing cells, as well as elevated GFAP expression, could be associated with elevated apoptosis, the mechanisms responsible for the overrepresentation of SATB2-expressing cells detected MAST1-mutant organoids remained unclear. As several patients with N-terminal MAST1 mutations have been diagnosed with the mega-corpus-callosum syndrome^24^ and because SATB2 regulates the specification of callosal projection neurons that send axons through the corpus callosum^45^, we investigated the establishment of corpus callosum-like neuronal projections between two organoids using a previously validated tissue-chip platform^46,47^. iCtrl or MAST1-mutant organoids were positioned in chambers connected by a microfluidic channel, and after 3 months, neuronal processes within the channel were analyzed (**Figure 1P**). While iCtrl organoids produced visible processes in the connecting channel, no obvious processes were observed in channels connecting MAST1-mutant organoids. Consistent with this observation, βIII-tubulin expression was significantly lower in protein lysates collected from channels connecting mutant organoids than in lysates from channels connecting iCtrl organoids (**Figure 1P**). These results demonstrate that overrepresentation of SATB2-expressing cells and impaired long-range neuronal projections paradoxically co-exist in MAST1-mutant organoids, suggesting positive effects of MAST1 mutations on the specification of SATB2-expressing cells and negative effects on neurite outgrowth in organoids. Collectively, our observations support the idea that the microcephaly-and corpus callosum-related growth deficits detected in MAST1-mutant organoids are associated with increased neuronal apoptosis, reduced neuropil, and changes in relative cell compositions.

### Organoids with MAST1 mutations show disrupted neuronal morphological development and elevated apoptosis before the emergence of microcephaly-associated growth deficits

Phenotypes detected in 5-month-old organoids are likely caused by both primary deficits directly induced by MAST1 mutations and secondary deficits, such as elevated neuronal apoptosis and chronic tissue stress. To gain insights into when and where the observed phenotypes originate, we investigated MAST1 expression at earlier developmental stages, including in PSCs, telencephalic neural progenitors of SNRs, and 1-month-old SNR-derived organoids comprising both telencephalic neural progenitors and neurons, using qRT-PCR (**Figures S6A and S6B**). MAST1 expression was undetectable in PSCs and SNR progenitors but increased approximately 20-fold in 1-month-old organoids. This increase correlated with an increase in the expression of the neuronal marker *TUBB3* (TUJ1), suggesting neuronal MAST1 expression. We validated preferential neuronal MAST1 expression using our previously reported single-cell RNA-sequencing (scRNA-seq) dataset obtained from 1-month-old SNR-derived organoids (**Figure S6C)** and datasets obtained from fetal human cortical tissue by others^48^. Together, these results demonstrate that MAST1 is preferentially expressed in nascent neurons early during human brain development. This is consistent with the results of a previous study of Mast1 expression in the mouse brain^24^.

Interestingly, although MAST1 is expressed in 1-month-old organoids, we observed no size differences between control and MAST1-mutant organoids at this time point (**Figures 2A and 2B**), suggesting that 1-month-old organoids could be used for studying the underlying pathogenic mechanisms disrupted by MAST1 mutations that later result in the microcephaly-associated size deficits observed in 5-month-old organoids. To gain insights into the pathogenic mechanisms, we first performed bulk RNA-seq on multiple control and patient organoids (**Figure S7**). Patient samples showed substantially different gene expression profiles compared with control samples and clustered separately from controls on the principal component analysis plot (**Figures S7A and S7B**), which could be related to different proportions of neurons and neural progenitors detected via bulk RNA-seq deconvolution analysis (**Figure S7C**). Despite these differences, however, both control and patient 1-month-old organoids transcriptionally recapitulated similar developmental stages, most resembling fetal human brain tissue at post-conception weeks 9–13 (**Figure S7D**), which is consistent with our previous characterization of 1-month-old organoids^49^. Analysis of upregulated differentially expressed genes in patient organoids revealed enrichment of Gene Ontology (GO) terms related to the cell cycle, DNA replication, and p53 signaling, with the following pathway-enriched genes: *MCM2–7*, *CDKN1A*, *CDK2*, *GADD45B*, and *TP53* (**Figures S7E–S7H and Table S3**). The downregulated genes, including *GAD1/2*, *GRM5*, *PTGER3*, and *GRIN2B*, largely fell into GO categories related to neuronal function, including neuroactive ligand–receptor interaction, calcium signaling, and long-term potentiation.

**Figure 2.**
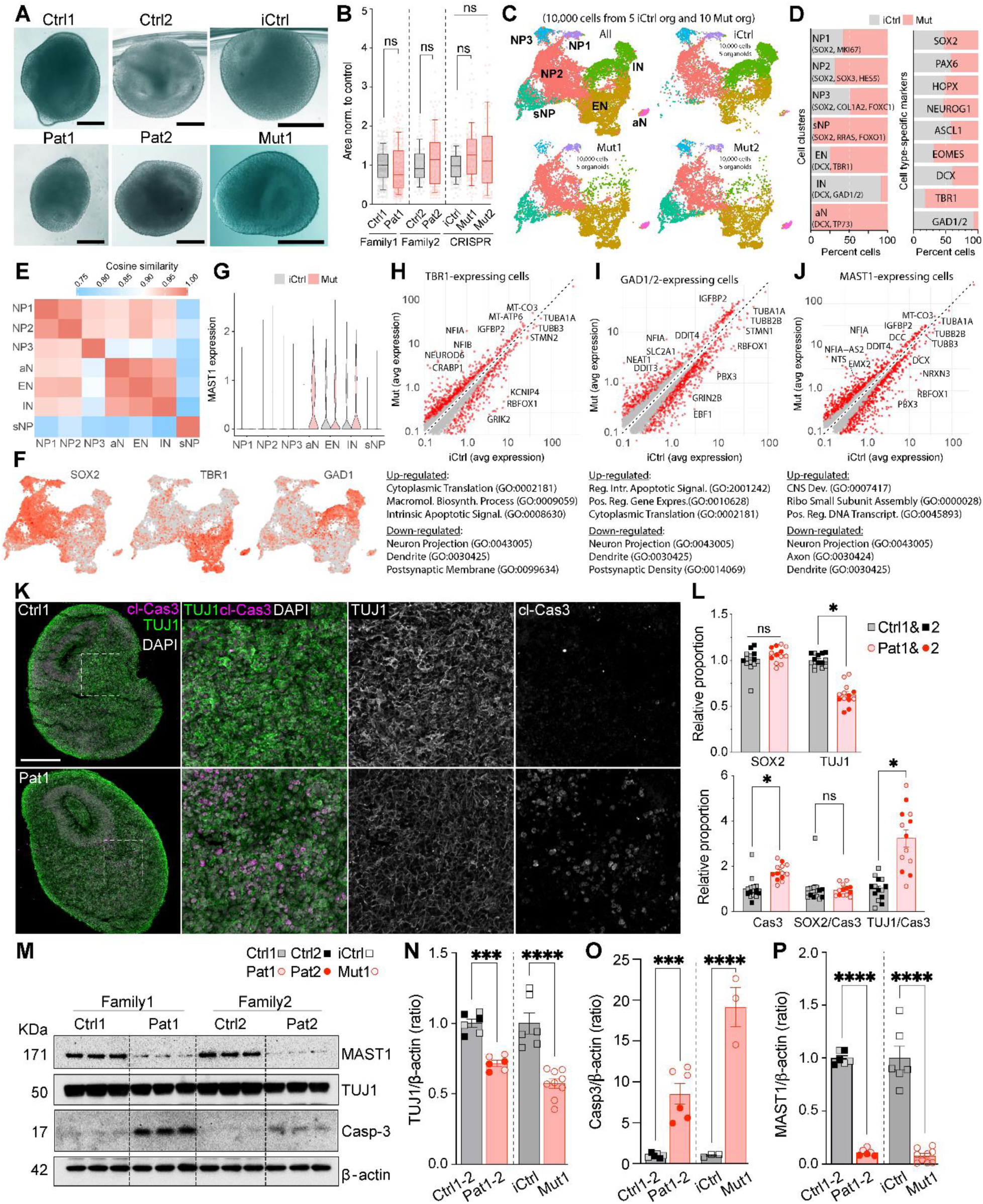
Disrupted morphological development and increased apoptosis in 1-month-old brain organoids with MAST1 mutations. (A) Images of 1-month-old SNR-derived brain organoids from control (ctrl1 and ctrl2), patient (pat1 and pat2), isogenic control (iCtrl), and MAST1-mutant (mut1) lines. Scale bar: 200 µm. (B) Organoid size quantifications (ctrl1: n = 222, ctrl2: n = 74, pat1: n = 224; pat2: n = 121, iCtrl: n=122, mut1: n=125, and mut2: n=97 organoids produced in 4-6 differentiation batches). (C) UMAP visualizations of cells and clusters obtained from 1-month-old iCtrl (n = 5), mut1 (n = 5), and mut2 (n = 5) organoids. Abbreviations: NP, neural progenitor; sNP, senescence-related neural progenitor; EN, excitatory neuron; IN, inhibitory neuron; aN, apoptotic neuron. (D) Percent cells in each cluster (left) and percent cells expressing cell type-specific marker (right) in iCtrl and mutant organoids. (E) Heatmap of pairwise cosine similarity between different cell clusters. (F) UMAP casting for SOX2, TBR1, and GAD1 expression in all organoids. (G) Violin plots of MAST1 expression across cell clusters in iCtrl and mut organoids. (H-J) Scatter plots of differentially expressed genes in TBR1-(H), GAD1/2-(I), and MAST1-(J) expressing cells from iCtrl and mut organoids. Selected Gene Ontology terms and accession numbers are depicted below. (K) Images of 1-month-old organoid sections immunostained for TUJ1 and cl-Cas3 with DAPI. Scale bar: 200 µm. (L) Normalized cell proportions in control and patient organoids (n = 5-9 images from 9 ctrl1, 9 pat1, 5 ctrl2 and 5 pat2 organoids produced 4 differentiation batches). Symbols show different batches and lines. (M) Immunoblots of TUJ1, cl-Cas3 and MAST1 in 1-month-old organoid lysates from control and patient organoids (3 organoids per sample). Loading control: β-actin. (N-P) Quantifications of normalized TUJ1, cl-Cas3 and MAST1 expression in control, patient, isogenic control and mutant organoids (n = 3-9 lysates/3 organoids per lysate sample, produced in 3 differentiation batches). Symbols show different batches and lines. Scale bars: 200 µm (A and K). Error bars = mean ± SEM. ns, not significant; *P<0.05, **P<0.01, ***P<0.001, and ****P<0.0001; unpaired two-tailed t-test.

To gain insights into cell type-specific effects of MAST1 mutations, we performed scRNA-seq on 1-month-old iCtrl and MAST1-mutant organoids (**Figures 2C–2J**). Consistent with bulk RNA-seq results, MAST1-mutant organoids showed increased proportions of cells in NP clusters, reduced proportions of cells in neuronal (excitatory neuron [EN] and inhibitory neuron [IN]) clusters, and a significantly elevated proportion of apoptotic neurons (aN) (**Figures 2C–2F**). Interestingly, despite an overall reduction in the proportion of neurons in MAST1-mutant organoids, these organoids showed elevated proportions of cells expressing EN marker *TBR1* and a reduced proportion of cells expressing IN markers *GAD1/2* (**Figure 2D**). However, both *TBR1*-and *GAD1/2*-expressing cells were characterized by elevated expression of apoptosis-related genes, including *TP73*, *DDIT3/4*, *CLU*, and *BBC3*, and reduced expression of genes associated with dendritic and axonal morphological development, including *TUBB3*, *STMN1/2*, and *DCX* (**Figures 2H and 2I**), suggesting the more rapid loss of INs may be due to apoptosis and that ENs may be more resilient or overproduced initially in MAST1-mutant organoids. The overproduction of ENs idea is supported by the increased proportions of *EOMES*-expressing intermediate progenitor cells (IPCs) detected in MAST1-mutant organoids compared with iCtrl organoids (**Figure 2D**). No major differences were observed in the proportions of cells expressing other neurogenesis markers, including *NEUROG1* and *ASCL1*, or NP markers, including *SOX2*, *PAX6*, and *HOPX* (**Figure 2D**).

Another interesting observation obtained using scRNA-seq is that the increased proportions of cells in NP clusters of MAST1-mutant organoids are largely attributable to the gain of a new NP cluster, characterized by the expression of cellular stress-or senescence-related genes, including *RRAS*, *FOXO1*, and *ITPR1* (sNP, **Figures 2C–2E**). These cells expressed no MAST1 (**Figure 2G**) and were largely absent in samples obtained from iCtrl organoids (**Figure 2C**), suggesting that these cells are likely specified as a result of non-cell-autonomous influences in MAST1-mutant organoids. Indeed, most *MAST1*-expressing cells were distributed to neuronal clusters (**Figure 2G**), and, similarly to *TBR1*-and *GAD1/2*-expressing neurons, they were characterized by increased expression of apoptotic genes and reduced expression of genes associated with dendritic and axonal morphological development in MAST1-mutant organoids (**Figure 2J**).

We validated the key observations from RNA-seq experiments via immunostaining and western blotting with antibodies against the NP marker SOX2, neuronal marker TUJ1, and the apoptosis marker cleaved caspase-3 in control, patient-derived, and engineered MAST1-mutant organoids (**Figures 2K–2P**). Indeed, MAST1-mutant organoids showed reduced TUJ1 and increased cleaved caspase-3 expression compared with control organoids. Interestingly, MAST1 protein expression was also dramatically reduced in MAST1-mutant organoids (**Figures 2M and 2P**), while no differences were detected in MAST1 mRNA expression levels by bulk RNA-seq or qRT-PCR or in the proportions of *MAST1*-expressing cells by scRNA-seq, suggesting that patient mutations substantially deplete MAST1 protein abundance in neurons, which may lead to impaired neuronal morphological development and apoptosis.

### Human neurons with MAST1 mutations exhitibit impaired morphological development

To directly investigate effects of MAST1 mutations on neuronal morphological development, we generated stem cell-derived cortical neurons from control, patient, and isogenic engineered MAST1-mutant lines following our established differentiation protocol^50^ (**Figure 3**). To visualize neuronal morphologies, two-dimensional cultures were immunostained with antibodies against axonal and dendritic microtubule protein TUJ1 and dendrite-specific microtubule-associated protein MAP2 (**Figure 3A**). Patient cultures consistently contained reduced proportions of TUJ1-expressing neurons (**Figure 3B**), and the neurons still generated in patient cultures exhibited significantly shorter dendrites with fewer branches (**Figures 3C and 3D**) as well as reduced TUJ1, MAP2, and MAST1 expression levels in both dendrites and axons (**Figures 3E–3I**). These results confirm that patient MAST1 mutations affect the expression levels of MAST1 and neuronal microtubule-associated proteins TUJ1 and MAP2, resulting in impaired neuronal morphological development.

**Figure 3.**
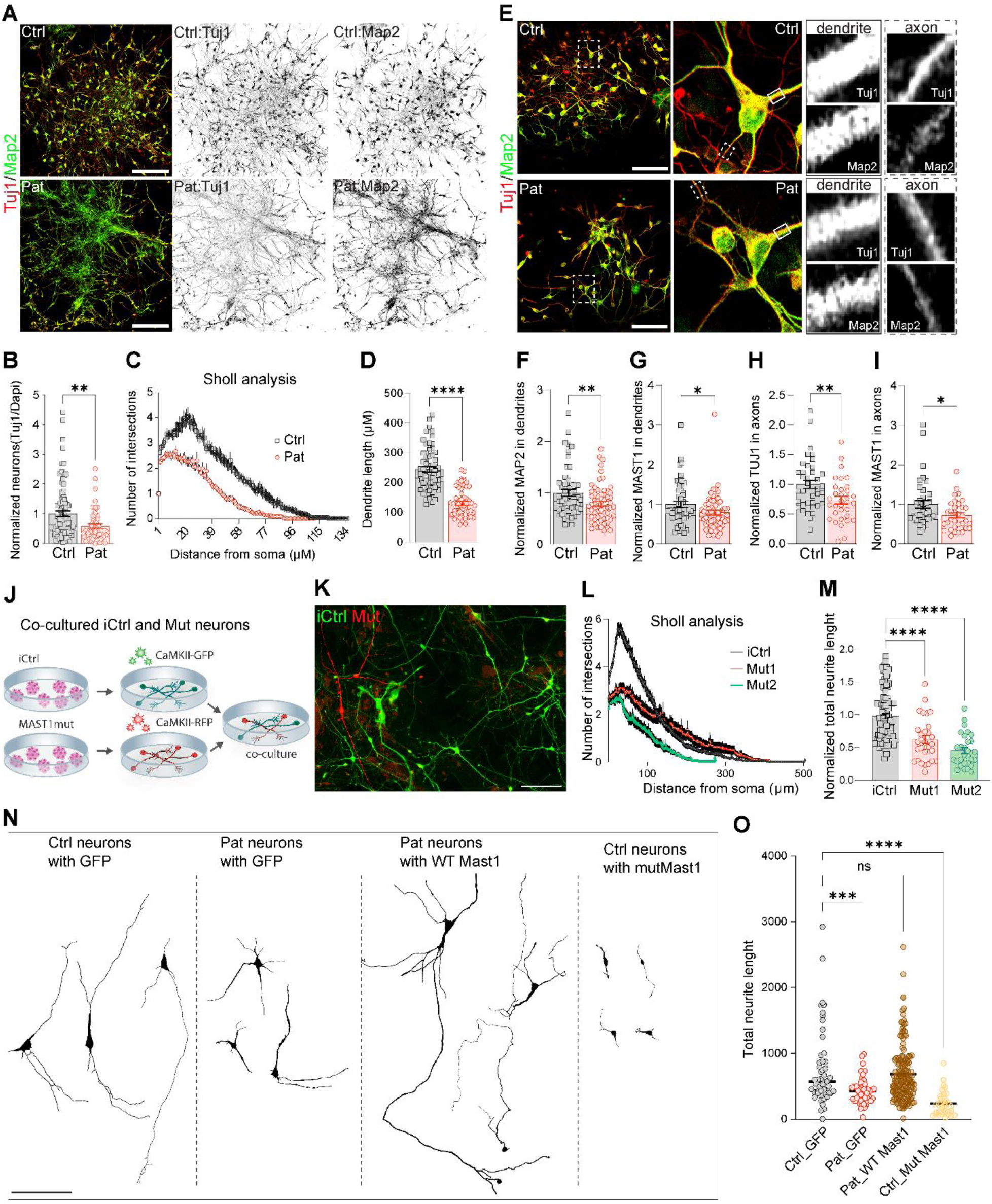
Altered neurite outgrowth from human iPSC-derived cortical neurons with MAST1 mutation. (A) Images of 3-week-old control and patient neuronal cultures immunostained for TUJ1 and MAP2. Scale bar: 200 µm. (B) Quantification of normalized proportions of TUJ1-expressing neurons (n = 52 and 52 images of 4 wells of ctrl1 and pat1 cultures obtained from in 2 differentiation batches). (C-D) Sholl analysis of MAP2-expressing dendrites (C) and quantification of total dendrite length (D) (n = 135 and 112 neurons collected from images of 4 wells of ctrl1 and pat1 cultures obtained in 2 differentiation batches). (E) Images control and patient neurons immunostained for TUJ1 and MAP2 with zoom-ins onto selected segments along dendrites (TUJ1+/MAP2+) and axons (TUJ1+/MAP2−). Scale bar: 100 µm. (F-I) Quantification of relative MAP2 (F) and MAST1 (G) and TUJ1 (H) and MAST1 (I) expression levels in dendrites (F–G) and axons (H-I) (n = 54 and 61 dendrites and 38 and 33 axons of 52 Ctrl1 and 52 Pat1 neurons collected from images of 4 wells of ctrl1 and pat1 cultures obtained in 2 differentiation batches). (J) Schematic depicting experimental design for co-culturing iCtrl and mutant neurons expressing CaMKII-GFP and CaMKII-RFP lentiviral reporters. (K) Images of cultured iCtrl and mutant neurons expressing CaMKII-GFP and CaMKII-RFP, respectively. Scale bar: 100 µm. (L and M) Sholl analysis of neurites (L) and quantification of normalized total neurite length (M) (n = 90 iCtrl, 30 mut1, and 30 mut2 neurons collected from images of 6 wells obtained in 3 differentiation batches) (N) Images of traced neurons from cultures of control and patient neurons expressing GFP (left) and patient and control neurons expressing WT-MAST1_T2A_GFP and mut-MAST1_T2A_GFP, respectively. Scale bar: 100 µm. (O) Quantification of normalized total neurite length (n = 51 GFP-expressing ctrl1, 47 GFP-expressing pat1, 156 WT-MAST1-expressing pat1, and 31 mut-MAST1-expressing ctrl1 neurons collected from images of 8 wells obtained in 4 differentiation batches). Scale bar: 200 (A) and 100 µm (E, K and N). Error bars = mean ± SEM. ns, not significant; *P<0.05, **P<0.01, ***P<0.001, and ****P<0.0001; two-tailed unpaired t-test (B), (D) and (F-I), ordinary one-way ANOVA test (M) and (O).

Impaired morphological development of patient neurons could result from MAST1 mutations, disrupted expression of other genes in patient neurons, and non-cell-autonomous effects induced by different neuronal density and altered expression of neuronal morphogens. To study the morphological deficits induced by MAST1 mutation, we generated cortical neurons from iCtrl and MAST1-mutant lines and transduced them with lentiviruses carrying different fluorescent proteins (green or red fluorescent protein [GFP or RFP], respectively or vice versa) under control of the neuron-specific CaMKIIα promoter to label more mature cortical neurons^50^. Upon dissociation, we co-cultured iCtrl and mutant neurons to ensure exposure to the same experimental conditions (**Figures 3J and 3K**). Consistent with the observations in patient neurons, MAST1-mutant neurons showed significantly impaired dendritic arbors compared with co-cultured iCtrl neurons (**Figures 3L and 3M**), indicating that MAST1 mutation is responsible for the morphological deficits in patient neurons.

Finally, to test whether expression of exogenous WT human MAST1 in patient neurons can compensate morphological deficits, we sparsely expressed WT MAST1 under the control of the doxycycline (DOX)-controllable promoter in separately cultured patient neurons using lentiviruses and an optimized DOX concentration (1 μg/μl) to reach MAST1 expression levels detected in control neurons. We used separately cultured control and patient neurons infected with GFP under the DOX-controllable promoter as positive and negative controls, respectively, in these experiments. Patient neurons expressing exogenous WT MAST1 showed significantly longer neurites than those expressing GFP only (**Figures 3N and 3O**). Interestingly, expression of mutated MAST1 in control neurons induced neurite outgrowth deficiency, suggesting that mutated MAST1 likely produces a dominant negative effect on neuronal morphological development.

### MAST1 is a mitochondria-associated protein that regulates mitochondrial abundance and distribution in developing human neurons

To gain further insights into the cellular and molecular mechanisms responsible for the detected deficits in neurons with MAST1 mutation, we examined the expression pattern of MAST1 in control 1-month-old organoids via immunostaining with anti-MAST1 antibodies (**Figure 4**). We observed a punctuated pattern of MAST1 expression in the neuronal layers at the periphery of organoid sections, which indicates potential protein localization to a specific subcellular compartment. Indeed, MAST1-containing puncta showed overlap with mitochondrial puncta visualized using a mitochondria marker heat shock protein 60 (HSP60) (**Figure 4A**), and no overlap was detected between MAST1 puncta and an endosomal marker early endosome antigen 1 (EEA1) or stress granules marker Ras GTPase-activating protein-binding protein 1 (G3BP1) (**Figure S8**), suggesting preferential MAST1 interaction with mitochondria. To validate this observation and investigate MAST1 distribution across neuronal soma and dendrites, we immunostained iPSC-derived control and mutant neuronal cultures with MAP2, MAST1, and another mitochondrial marker TOM20 (**Figures 4B and 4C**). We also isolated mitochondria from iCtrl and MAST1-mutant organoids to investigate MAST1 expression levels specifically in the mitochondrial protein lysates via western blotting (**Figure 4D**). We observed partial co-localization of MAST1 and TOM20 in both the cell soma and dendrites of iCtrl and MAST1 mutant neurons and significantly reduced total mitochondrial volume at the cell soma and mitochondria density in dendrites of MAST1-mutant neurons compared with iCtrl neurons (**Figures 4B and 4C**). We also found that relative MAST1 expression was significantly reduced in the mitochondrial fraction of MAST1-mutant organoids as compared with iCtrl organoids (**Figure 4D**), suggesting the patient mutation affects MAST1 association with mitochondria. These results indicate that MAST1 associates with neuronal mitochondria, regulating mitochondria abundance and distribution in developing human neurons, and that the patient MAST1 mutations disrupt this association to reduce neuronal mitochondria abundance and their presence in dendrites.

**Figure 4.**
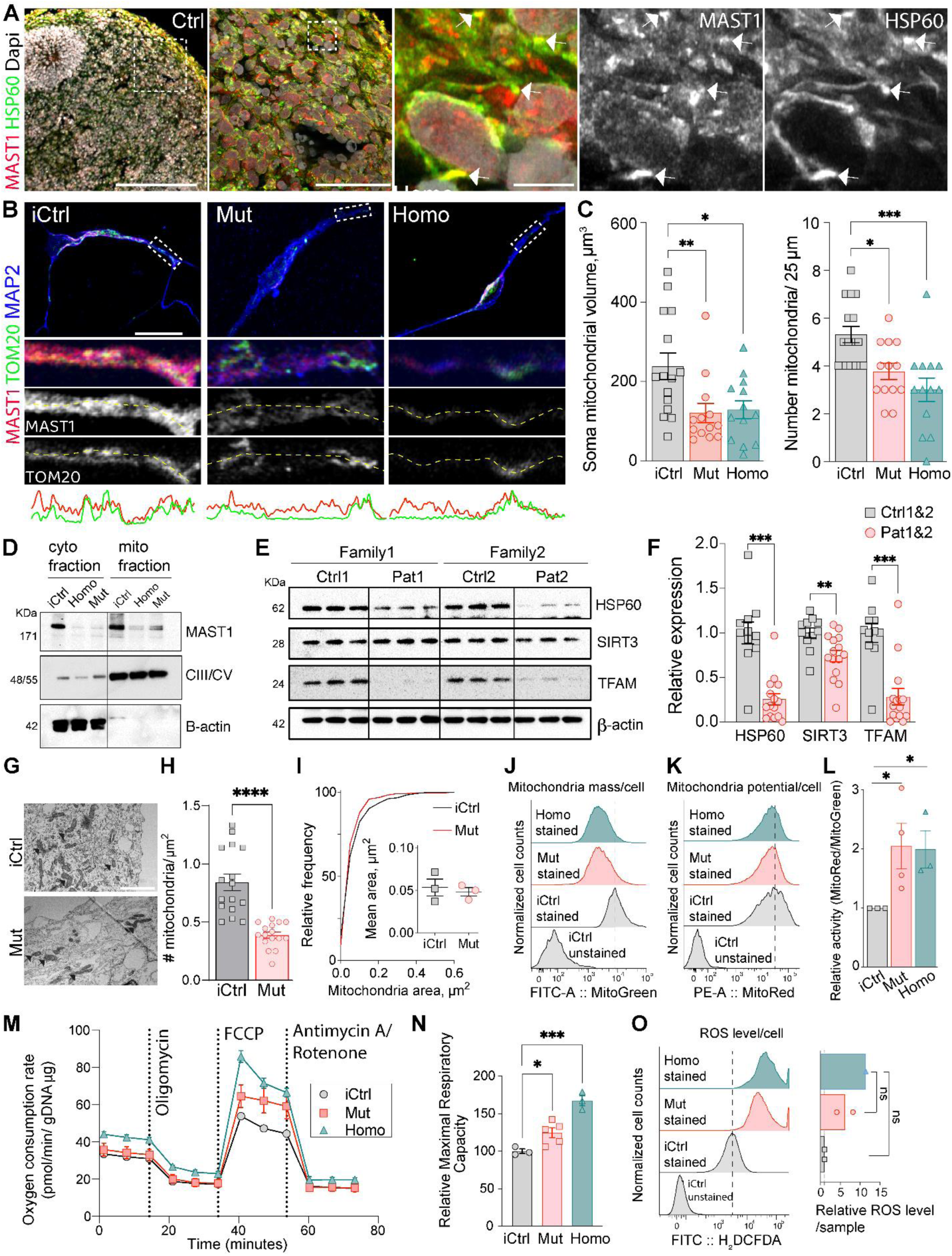
Association of MAST1 with neuronal mitochondria and of MAST1 mutations with mitochondrial deficits. (A) Images of 1-month-old control organoid sections immunostained for HSP60, MAST1, and DAPI. Arrowheads indicate puncta with MAST1 and HSP60 colocalization. Scale bars: 200, 50, and 5 µm. (B) Images of control and mutant neurons immunostained for MAST1, TOM20 and MAP2 and expression profiles of MAST1 (red) and TOM20 (green) in dendrites. Scale bar: 20 µm. (C) Quantification of somatic mitochondrial volume (left) and mitochondrial quantity in dendrites (right) (n= 15 iCtrl, 13 mut, and 13 homo mut neurons collected from images of 9 wells obtained in 3 differentiation batches). (D) Immunoblots of MAST1, mitochondrial complex III/V, and β-actin in cytosolic and mitochondrial lysates isolated from 1-month-old iCtrl and mutant organoids. (n=14 organoids per group from 3 differentiation batches). Loading controls for cytosolic and mitochondrial lysates are β-actin and complex III/V, respectively. (E) Immunoblots of HSP60, SIRT3 and TFAM in protein lysates from 1-month-old control and and patient organoids (3 organoids per sample obtained in 2-3 batches). Loading control: β-actin. (F) Quantification of normalized HSP60, SIRT3 and TFAM expression (n= 6 ctrl1, 5 ctrl2, 8 pat1, and 7 pat2 lysates/3 organoids per lysate obtained in 2-3 differentiation batches). (G) Representative images of mitochondria at the periphery of 1-month-old iCtrl and mutant organoid sections. Scale bar: 2 µm. (H-I) Quantification of mitochondria density (H) and size (I) (n=16 and 16 images collected from 3 iCtrl and 3 mut organoids obtained in 2 differentiation batches). (J-L) Flow-cytometry histograms of mitochondrial mass (MitoGreen, J), membrane potential (MitoRed, K) and relative mitochondrial activity (MitoRed/MitoGreen, L) in cells obtained from 1-month-old iCtrl and mutant organoids; Negative control: unstained cells from iCtrl organoids (n=14 organoids per group from 3-4 differentiation batches). (M-N) Quantification of oxygen consumption rate (M) and maximal respiratory capacity (N) using Seahorse stress test in cultures of 3-4-week-old iCtrl and mutant neurons normalized to gDNA collected from corresponding cultures. (n=3 iCtrl, 5 mut, and 3 homo mut wells obtained in 3 differentiation batches). (O) Flow-cytometry histograms of ROS levels measured by H2DCFDA staining in cells obtained from cultures of iCtrl and mutant neurons; Negative control: unstained cells from iCtrl organoids (n=2 cultures from 3 differentiation batches) Scale bar: 200, 50 (zoom-in 1), and 5 (zoom-in 2) (A), 20 (B), and 2 µm (G). Error bars = mean ± SEM. *P<0.05, **P<0.01, ***P<0.001, and ****P<0.0001; ordinary one-way ANOVA test (C), (L), (N) and (O) unpaired two-tailed t-test (F) and (H).

### MAST1-mutant neurons show mitochondrial deficits

To confirm the effect of MAST1 mutation on mitochondrial abundance in brain organoids, we measured the expression levels of key mitochondrial marker proteins that work in different pathways, including HSP60, NAD-dependent protein deacetylase sirtuin-3 (SIRT3), and transcription factor A, mitochondrial (TFAM), in protein lysates obtained from control and patient organoids by western blotting (**Figure 4E**). All proteins showed significantly reduced expression levels in MAST1-mutant organoids (**Figure 4F**). Moreover, we observed a significant reduction in mitochondrial count, but not size, in MAST1-mutant organoid sections on transmission electron microscopy (**Figures 4G–I**). Collectively, these results confirm diminished mitochondrial abundance in MAST1-mutant brain organoids, which is consistent with the idea that MAST1 is an important regulator of mitochondrial abundance in human neurons during development.

To determine whether impaired mitochondrial abundance in MAST1-mutant neurons is also associated with altered mitochondrial function, we co-stained cells obtained from iCtrl and mutant organoids with cell permeable mitochondrial dyes MitoTracker-Red and MitoTracker-Green that accumulate the mitochondria in membrane potential-dependent and -independent manners, providing cellular fluorescent signals proportional to the cellular mitochondrial mass only and the mitochondrial mass plus potential, respectively (**Figures 4J and 4K**). Again, significantly lower MitoTracker-Green signal was detected in MAST1-mutant samples by fluorescence-activated cell sorting (FACS), consistent with reduced mitochondrial abundance, but no differences were found in MitoTracker-Red signal between iCtrl and mutant cells, suggesting that mitochondria in mutant cells are more hyperpolarized compared with those in iCtrl cells. Indeed, the ratio of MitoTracker-Red to MitoTracker-Green signals was significantly greater in mutant cells than in iCtrl cells (**Figure 4L**), suggesting an increased proportion of hyperpolarized and potentially hyperactive mitochondria in MAST1-mutant cells.

To directly investigate neuronal mitochondrial activity in iCtrl and MAST1-mutant neurons, we measured the oxygen consumption rates in cultured iCtrl and MAST1-mutant cortical neurons using a Seahorse mitochondrial stress test assay (**Figure 4M**). Mutant neurons demonstrated significantly elevated FCCP-stimulated maximal oxygen consumption, suggesting the remaining mitochondria have an increased intrinsic respiratory capacity (**Figure 4N**). A side effect of mitochondrial hyperactivity is the excessive production of unstable ROS, which can cause cell membrane and DNA damage and trigger apoptosis^51^. As elevated apoptosis was detected in MAST1-mutant neurons, we measured ROS production in cells dissociated from iCtrl and MAST1-mutant organoids via staining with a cell permeable ROS-detection probe H_2_DCFDA (2′,7′-dichlorodihydrofluorescein diacetate) and FACS (**Figure 4O**). MAST1-mutant cells demonstrated approximately eight-fold elevated ROS production compared with control cells, indicating that elevated mitochondrial activity in MAST1-mutant neurons results in increased ROS production, which may be responsible for neuronal apoptosis.

### MAST1 mutations disrupt mitochondrial stabilization in neurites

We reasoned that MAST1 may regulate mitochondrial trafficking as it directly binds microtubules^25^, which are essential for mitochondrial transportation in neurons, and we observed reduced mitochondrial density in the dendrites of MAST1-mutant neurons compared with that in controls (**Figures 4B and 4C**). To examine neuronal mitochondrial trafficking, we generated a lentiviral reporter that carries RFP under control of the CaMKIIα promoter and the T2A-linked mitochondria-targeted Emerald GFP (mito-Emerald) for mitochondrial visualization in RFP-expressing neurons. iCtrl and MAST1-mutant iPSC-derived cortical neurons were infected with this reporter. We then recorded mitochondrial locations in neurites of iCtrl and MAST1-mutant neurons over time using live-cell imaging (**Figures 5A and 5B**). The mitochondrial density was approximately 30% lower in neurites of MAST1-mutant neurons than in those of control neurons (**Figure 5C**), which is consistent with our previous observations (**Figure 4**). Interestingly, we also found that MAST1-mutant neurites contained approximately 30% fewer stationary mitochondria compared with control neurites (**Figure 5D**) and that the speed of forward movement of mobile mitochondria was approximately 80% faster for MAST1-mutant neurites than for control neurites (**Figure 5E**). No differences were detected in the speed of the backward movement or the length of mitochondrial displacement (**Figures 5F–5H**). These results suggest that MAST1 is specifically involved in stabilizing anterogradely moving mitochondria in neurites and that MAST1 mutations disrupt this stabilization, leading to an increased proportion of mobile and unstable mitochondria.

**Figure 5.**
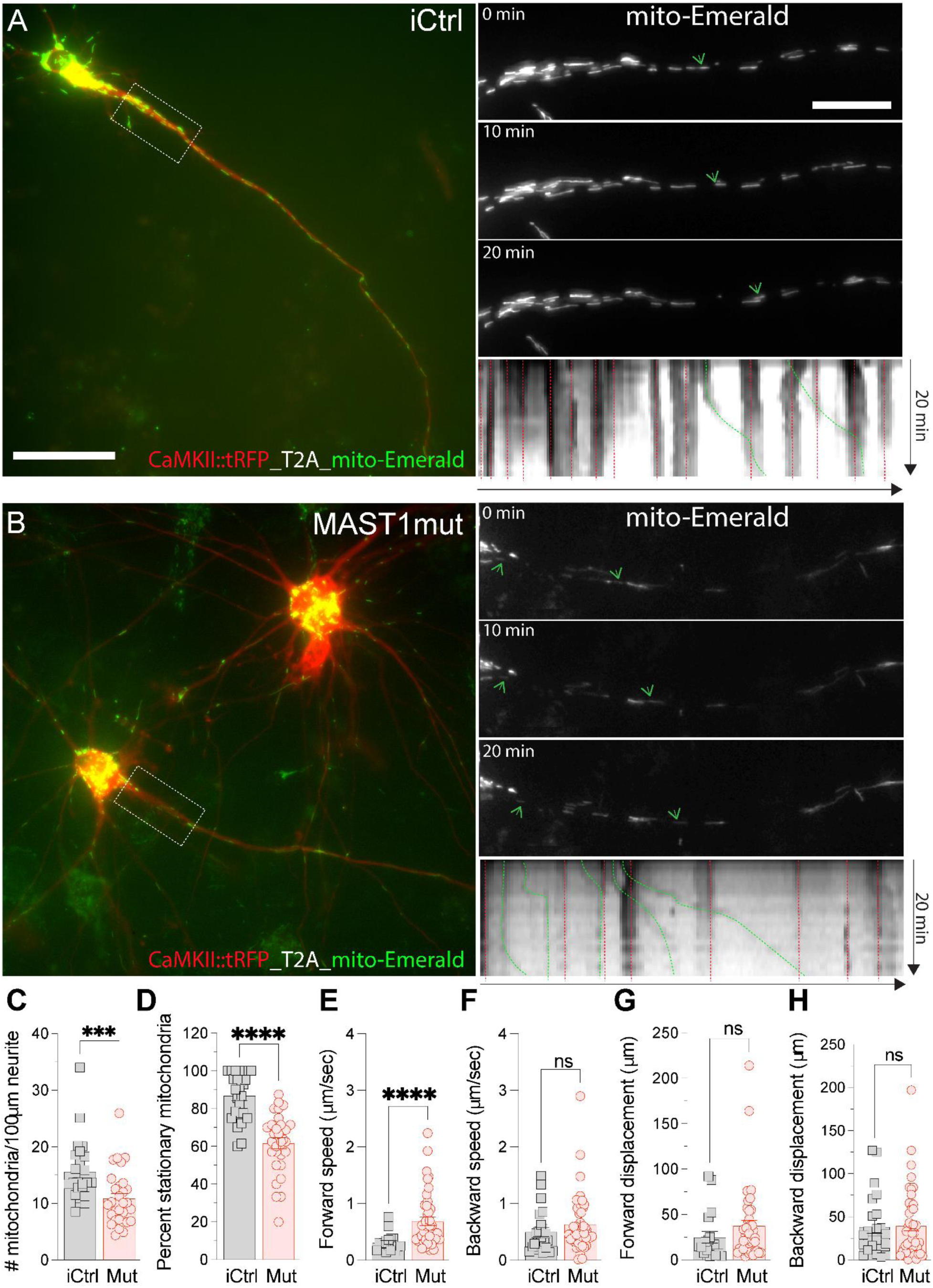
Disrupted mitochondrial trafficking in human cortical iPSC-derived neurons with MAST1 mutations. (A-B) Images of iCtrl (A) and mutant (B) neurons expressing CaMKII::tRFP_T2A_mito-Emerald reporter (left) and images of selected dendrites (right) obtained at 0 (top), 10 (middle), and 20 (bottom) minutes during live-cell imaging. Arrowheads indicate individual mitochondria. Scale bars: 50 µm (left) and 20 µm (right). (C-H) Quantification of mitochondria density (C), percent stationary mitochondria (D), forward (E) and backward (F) speed (µm/s) and forward (G) and backward (H) displacement (µm) in iCtrl and mut neurites (n=25 iCtrl and 30 mut neurites collected from neurons in 4 differentiation batches). Error bars = mean ± SEM. ns, not significant; *P<0.05, **P<0.01, ***P<0.001, and ****P<0.0001; unpaired two-tailed t-test.

### Creatine and NAC treatments compensate neuronal deficits caused by MAST1 mutations

The detection of reduced mitochondrial stabilization and abundance in MAST1-mutant neurons along with elevated activity and increased ROS production suggest that disrupted energy distribution rather than production may be responsible for the deficits detected in MAST1-mutant neurons. To test whether improved energy distribution and ROS inhibition can eliminate the deficits in MAST1-mutant organoids, we treated them with 4 mM creatine or 2 mM NAC (**Figure 6**). Creatine facilitates ATP delivery to neurites through the creatine kinase pathway^52^, and NAC suppresses ROS accumulation through glutathione synthesis^53^. First, we measured organoid size and the ROS levels in cells obtained from untreated and treated organoids. For size evaluation, we selected the 2-month time point, as it was the earliest time point at which size deficits were detected (**Figure S9**). After 7 weeks of treatment, we measured organoid size under bright-field imaging (**Figures 6A and 6B**) and ROS levels using H_2_DCFDA staining and FACS (**Figure 6C**). We found that MAST1-mutant organoids, produced from both patient and engineered lines, showed significantly greater size if treated with creatine and NAC (**Figure 6B**); however, the treated organoids did not reach the size of control untreated organoids, suggesting that the treatments compensated, but not completely rescued, the size deficits in organoids. On the other hand, ROS levels were significantly lower in treated MAST1-mutant cells than in untreated cells and approached the ROS levels detected in cells from control organoids (**Figure 6C**), suggesting rescue of this effect of MAST1 mutation.

**Figure 6.**
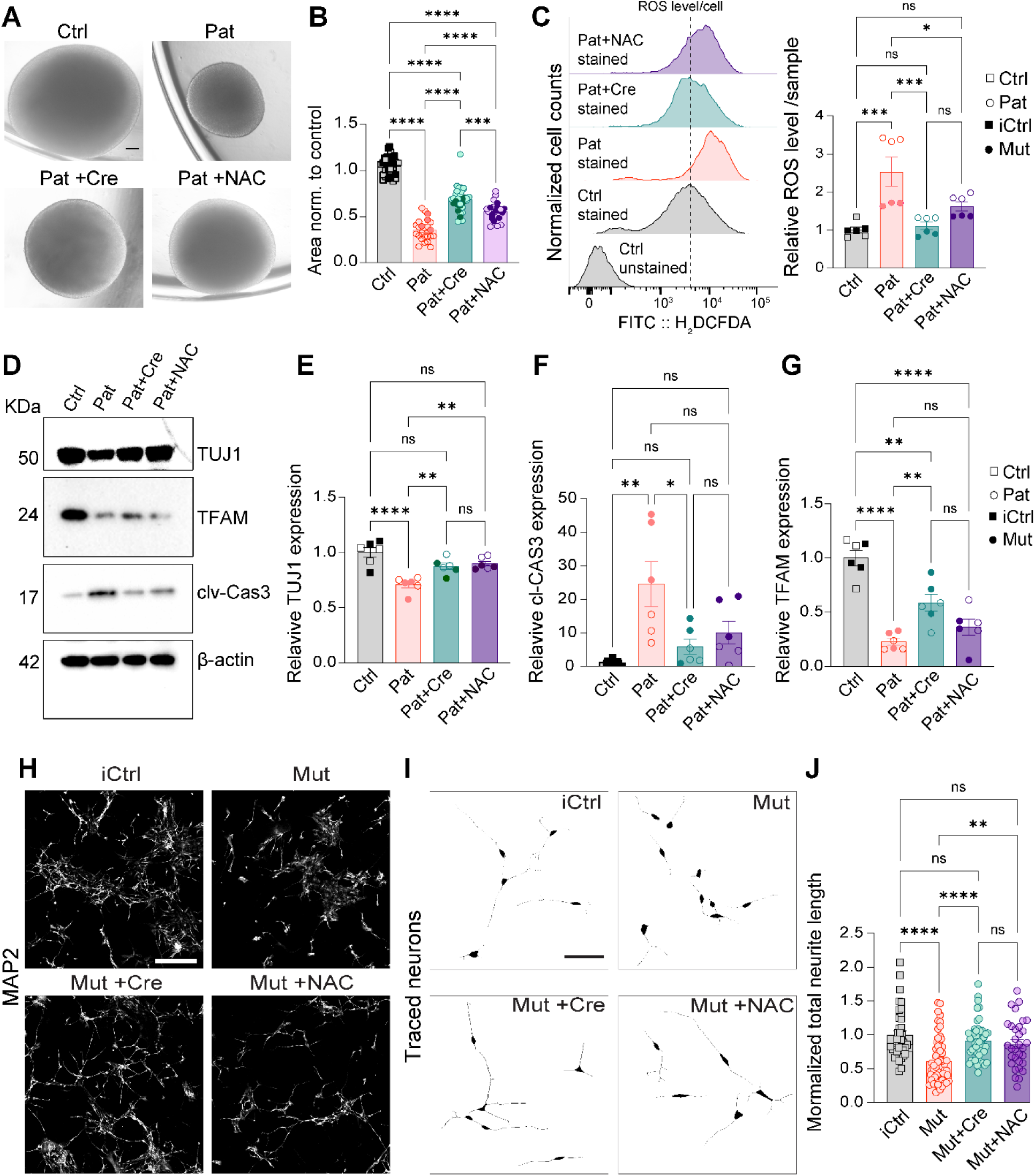
Amelioration of neuronal deficits in MAST1-mutant organoids and neurons with prolonged creatine and NAC treatment. (A) Images of 2-month-old control (ctrl1), patient (pat1) and pat1 organoids treated with creatine (pat1+Cre) or *N*-acetyl-L-cysteine (pat1+NAC). Scale bar: 200 µm. (B) Quantification of normalized sizes of 2-month-old ctrl1, pat1, iCtrl, and mut organoids (n= 15 ctrl1, 20 pat1, 20 pat1+Cre, 10 pat1+NAC, 17 iCtrl, 4 mut, 14 mut+Cre, and 12 mut+NAC organoids obtained from 8 differentiation batches). (C) Flow-cytometry histograms of ROS levels measured by H2DCFDA staining in cells obtained from 2-month-old ctrl1, pat1, iCtrl, and pat1/mut organoids treated with creatine or *N*-acetyl-L-cysteine (n = 3 ctrl1, 3 pat1, 3 pat1+Cre, 3 pat1+NAC, 3 iCtrl, 3 mut, 3 mut+Cre, and 3 mut+NAC samples/3 organoids per sample obtained in 8 differentiation batches). (D) Immunoblots of TUJ1, cl-Cas3, and TFAM in protein lysates from 2-month-old ctrl1, pat1, and pat1 organoids treated with creatine (pat1+Cre) or *N*-acetyl-L-cysteine (pat+NAC). Loading control: β-actin. (E-G) Quantification of normalized TUJ1 (E), cl-Cas3 (F), and TFAM (G) expression (n= 3 ctrl1, 3 pat1, 3 pat1+Cre, 3 pat1+NAC, 3 iCtrl, 3 mut, 3 mut+Cre, and 3 mut+NAC lysates/3 organoids per lysate obtained in 4 differentiation batches). (H-I) Images (H) and traces (I) of iCtrl, mut, and mut neurons treated with creatine (mut+Cre) or *N*-acetyl-L-cysteine (mut+NAC) immunostained for MAP2. Scale bar: 200 µm (left) and 50 µm (right). (J) Quantification of normalized total neurite length (n = 40 iCtrl, 60 mut, 49 mut+Cre, 32 mut+NAC neurons from 2 cultures obtained in 4 differentiation batches). Error bars show mean ± standard error of the mean (SEM). ns, not significant; *P<0.05, **P<0.01, ***P<0.001, and ****P<0.0001; ordinary one-way ANOVA test.

Because elevated ROS levels in MAST1-mutant cells could be caused by elevated mitochondrial activity due to reduced mitochondrial abundance and could lead to neuronal apoptosis, we next examined the expression levels of a mitochondrial marker TFAM to assess mitochondrial abundance as well as expression of the neuronal marker β3 tubulin (TUJ1) and apoptotic marker cleaved caspase-3 to evaluate neuronal apoptosis (**Figures 6D–6G**). Treated MAST1-mutant organoids demonstrated significantly increased TUJ1 expression (**Figure 6E**), reduced cleaved caspase-3 expression (**Figure 6F**), and moderately increased TFAM expression with creatine treatment, but not with NAC treatment (**Figure 6G**), suggesting that the treatments primarily alleviate downstream consequences of reduced mitochondrial abundance rather than directly correcting the mitochondrial deficit itself.

Finally, to observed whether the tested treatments can improve neurite outgrowth, we treated engineered MAST1-mutant cortical neurons with 4 mM creatine or 2 mM NAC for 3–4 weeks and measured dendrite outgrowth via immunostaining of MAP2 (**Figure 6H**). We found that treated MAST1-mutant neurons exhibited neurites that were similar in length than those of iCtrl untreated neurons and significantly longer than those of untreated MAST1-mutant neurons (**Figures 6I and 6J**), supporting the idea that impaired energy availability and oxidative stress are likely responsible for impaired neurite outgrowth from MAST1-mutant neurons.

## Discussion

The study identifies a neuron-specific protein kinase MAST1 as an important regulator of mitochondrial trafficking in human neurons. It also describes the cellular and molecular deficits detected in cortical neurons and telencephalic brain organoids generated from patients and engineered stem cells with patient-specific *MAST1* mutations. The detection of *MAST1* mutations in multiple individuals with brain malformations, severe intellectual disability, and neurodevelopmental abnormalities suggests that disrupted mitochondrial trafficking and associated mitochondrial and neuronal deficits may contribute to the neurological problems in these patients. Importantly, our results demonstrate that neuronal deficits in MAST1-deficient neurons can be largely compensated by prolonged treatment with creatine and NAC, which are two widely used compounds for energy, metabolism, and antioxidant support. Collectively, this study describes the cellular and molecular deficits induced by patient-specific *MAST1* mutations in human neurons and brain organoids, identifies disrupted mitochondrial trafficking as a candidate pathogenic mechanism, and provides new insights into potential therapeutic strategies for patients with *MAST1* mutations and other neurological disorders associated with mitochondrial dysfunction.

### MAST1-deficient human brain organoids and neurons exhibit cellular deficits

We observed growth deficits and alterations in cellular composition in 5-month-old MAST1-mutant organoids. Specifically, mutant organoids showed significantly elevated proportions of apoptotic MAP2-expressing neurons and reduced proportions of HOPX-expressing progenitors, CTIP2-expressing neurons, GABA-expressing inhibitory neurons, and S100b-expressing astroglia cells. On the other hand, we observed no effects on the proportions of TBR1-expressing deep layer cortical neurons and surprisingly elevated proportions of SATB2-expressing superficial layer cortical neurons as well as GFAP-expressing glial cells. These results are difficult to interpret as they likely reflect the cumulative effects of multiple direct and indirect (chronic and compensatory) actions of MAST1 mutations on different cell types during organoid development. To gain insights into the pathogenic mechanisms, we investigated the properties of cells in organoids at an early developmental time point, 1 month, when MAST1 expression is already detected but no differences in organoid growth are observed. In these organoids, which are comparable to human fetal cortical development at 9–13 weeks post-conception in terms of gene expression profiles, MAST1 expression was largely restricted to neurons, suggesting that nascent neurons are likely the earliest cells affected by MAST1 mutations. Indeed, we detected elevated expression of stress-and apoptosis-related genes and impaired expression of genes involved in regulating neuronal morphological development in nascent neurons from 1-month-old organoids with MAST1 mutations compared with iCtrl organoids. We validated apoptosis and morphological deficits in 1-month-old organoids via immunofluorescence and western blotting. We also observed significantly reduced proportions of MAP2-expressing neurons with impaired neurites in MAST1-mutant cultures compared with control cultures, confirming the roles of MAST1 in regulating neuronal survival and morphological development. The elevated proportions of NPs with increased expression of senescence markers, also detected in 1-month-old organoids, are likely secondary effects of increased neuronal apoptosis and stress induced by MAST1 mutation in neurons, as NPs showed no or limited MAST1 expression.

### MAST1-mutant neurons show reduced mitochondrial abundance and mitochondrial hyperactivity

We discovered MAST1 association with neuronal mitochondria and their co-localization at certain locations within the cell soma and neurites, suggesting a dynamic interaction between MAST1 and neuronal mitochondria or MAST1 association with specific spatially and functionally distinct mitochondrial populations^3^. To examine the impact of MAST1 mutation on mitochondrial properties in neurons, we measured mitochondrial abundance and activity in Ctrl and mutant neurons and organoids. Mutant neurons showed reduced mitochondrial abundance but significantly elevated activity, suggesting that the remaining mitochondria in MAST1-mutant neurons are hyperactive, working to compensate for their impaired abundance. Consistent with increased mitochondrial activity, we also detected significantly elevated ROS production in MAST1-mutant neurons. These results indicate that mitochondrial energy production is not impaired by MAST1 deficiency and suggest that altered mitochondrial distribution in neurites is responsible for the detected neuronal deficits.

### MAST1 mutation disrupts mitochondrial stabilization in human neurons

Mitochondrial distribution is tightly regulated in neurons, as localized energy production in distal compartments supports neurite outgrowth, action-potential generation and propagation, synaptic transmission, and plasticity^54^. Several motor, adapter, and stabilizing proteins have been identified as important regulators of mitochondrial trafficking and localization in axons and dendrites^6^. Notably, mitochondrial distribution may be differentially controlled in neurites of human neurons versus rodent neurons, as the human neurons have substantially longer axons and dendrites ^21,22^. Human neurons also exhibit distinct biophysical properties^55,56^, mitochondrial metabolism^23^, and exceptional longevity^57^, suggesting potentially unique requirements for the regulation of mitochondrial distribution, retention, and function in human neurons. The present study demonstrates that human neurons with mutated MAST1 have significantly reduced proportions of stationary mitochondria in neurites compared with control neurons, suggesting that MAST1 is involved in regulating mitochondrial retention in neurites. Interestingly, reduced mitochondrial stability in neurites has been associated with impaired neurite outgrowth and branching^12,14^, as also observed in MAST1-mutant neurons. The current understanding of how MAST1 exerts its action on mitochondrial stability and how this translates into neurite outgrowth remains incomplete. A potential mechanistic model may involve MAST1 residing on neuronal microtubules and dynamically interacting with certain transported mitochondria to facilitate their docking at specific locations. This model is supported by the previous observations of MAST1 interaction with microtubules^24,25^ and our observations of MAST1 expression in the mitochondrial fraction and its co-localization with neuronal mitochondria in neurites. Other possible mechanisms may involve MAST1 phosphorylation and association with other proteins involved in regulating mitochondrial trafficking, biogenesis, activity, dynamics, or mitophagy. Interestingly, lethal loss of *Drosophila* MAST kinase, *dop*, impairs dynein-dependent trafficking of lipid droplets by indirectly reducing dynein intermediate chain phosphorylation^37^, suggesting that MAST1 may regulate the dynein-dependent retrograde mitochondrial trafficking in neurites^7^. However, we observed no retrograde mitochondrial trafficking deficits in MAST1-mutant neurons, and no lethality rescue was achieved upon MAST1 expression in *Drosophila* lacking *dop*. Additional studies are required to investigate MAST1 interactions with other mitochondrial proteins.

### Creatine and NAC treatments ameliorate neuronal deficits in MAST1-mutant neurons

Prolonged treatment with creatine or NAC largely compensates the neurite outgrowth and apoptosis deficits detected in MAST1-mutant neurons, suggesting that these widely used compounds may be promising therapeutic agents for treating deficits in patients with MAST1 mutations and mitochondrial abnormalities. Creatine acts by entering cells through the SLC6A8 transporter and serving as a substrate for creatine kinase to produce diffusible energy buffer phosphocreatine^58^. Phosphocreatine can diffuse into axons and dendrites and facilitate local ATP regeneration at distal locations, supporting neurite outgrowth^52,59^. Thus, the effects of creatine treatment on MAST1-mutant neurons could be mediated by improved ATP buffering and regeneration in neurites. NAC is a redox modulator that mitigates mitochondrial oxidative stress and limits intracellular ROS accumulation^53^. The compensatory effects of NAC treatment on MAST1-mutant neurons are likely mediated by reduced mitochondrial stress and improved neuronal health and survival. Indeed, previous studies demonstrated that NAC can promote neurite extension and neuron survival under both normal and disease conditions^60,61^.

### Potential impact and limitations

The study connects disease-associated MAST1 variants with altered mitochondrial trafficking and neuronal morphological development in human models and identifies bioenergetic and redox pathways as potential points for therapeutic intervention. The limitations of this study include (1) the small number of affected individuals and patient-derived iPSC lines with specific *MAST1* variants; (2) the limited characterization of longitudinal and later phenotypes in MAST1-mutant organoids, as the study focused largely on the discovery of early cellular and molecular pathogenic mechanisms; and (3) the limited range of creatine and NAC concentrations selected based on previous reports^61,62^. Remaining questions are related to whether and how the C-terminal *MAST1* variants affect MAST1 kinase activity and phosphorylation of downstream substrates, how MAST1 interacts with mitochondria, and how MAST1 contributes to mitochondrial stabilization in neurites. It also remains unclear whether impaired mitochondrial stabilization is a cause or consequence of reduced mitochondrial abundance and elevated mitochondrial activity. Additional studies are needed to address these questions.

## Materials and Methods

### Patient sample collection and exome sequencing

Three unrelated patients with overlapping neurological phenotypes were evaluated by trio-based exome sequencing. All three patients had normal chromosomal microarray results prior to exome testing. Clinical exome sequencing was performed on genomic DNA from each patient and their unaffected parents, with written informed consent obtained from their legal guardians. Genomic DNA was extracted from peripheral blood, and parental sequencing data were used to identify de novo variants; biological parentage was confirmed through analysis of numerous Mendelian consistencies within each trio. Sequencing reads were aligned to the human genome reference (GRCh37), and identified de novo variants were confirmed by targeted Sanger sequencing. A literature review was conducted to identify additional individuals with de novo *MAST1* variants. Variant filtering and interpretation incorporated multiple population and disease databases, including dbSNP (build 151; http://www.ncbi.nlm.nih.gov/SNP/), the Genome Aggregation Database (gnomAD v4.1.0; http://gnomad.broadinstitute.org), the Human Gene Mutation Database (HGMD Professional 2025.4, QIAGEN Bioinformatics), and the ClinVar Database (https://www.ncbi.nlm.nih.gov/clinvar/).

### Variant identification and interpretation

Exome sequencing identified three de novo missense variants in the *MAST1* gene in three patients (**Table S2**). Two patients carried variants affecting the same amino acid, Pro1177, specifically *c.3530C>T (p.Pro1177Leu)* in Patient 1 and *c.3529C>T (p.Pro1177Ser)* in Patient 2, while Patient 3 carried a different de novo missense variant, *c.3889G>T (p.Asp1297Tyr)*. All three variants are absent from population variant databases, and the altered residues (Pro1177 and Asp1297) are highly conserved. The REVEL score for *c.3530C>T (p.Pro1177Leu)* is 0.50, and this variant has been reported as a disease-causing change in a patient with developmental disabilities (PMID: 31721002). The REVEL scores for *c.3529C>T (p.Pro1177Ser)* and *c.3889G>T (p.Asp1297Tyr)* are 0.32 and 0.26, respectively. Mega-corpus-callosum syndrome with cerebellar hypoplasia and cortical malformations (MIM #618273) is believed to result from a gain-of-function mechanism, with most pathogenic variants being de novo missense changes in *MAST1*. Taken together, the de novo status, evolutionary conservation, absence from population databases, and alignment with the known disease mechanism provide sufficient evidence to support that these three missense variants are likely pathogenic or pathogenic and are consistent with causing Mega-corpus-callosum syndrome with cerebellar hypoplasia and cortical malformations.

### Generation of iPSCs and CRISPR/Cas9 engineered lines with *MAST1* mutations

This study was approved by the Institutional Review Board at the University of Utah (IRB_00007551). Blood samples were collected from two unaffected parents and two patients. iPSCs were generated from peripheral blood mononuclear cells (PBMCs) using CytoTune-iPS 2.0 Sendai reprogramming kit. MAST1-mutant stem cell lines were generated using a CRISPR/Cas9 approach from the H9 line. We generated isogenic MAST1-engineered control stem cells with patient 1 (Asp1297Tyr)-specific heterozygous and homozygous mutations. Upon electroporation of 5 µM CRISPR/Cas9 plasmids into ∼3×10^6^ cells using the Amaxa Nucleofector System, cells were resuspended in E8 medium with ROCK inhibitor. Starting the next day, the cells were exposed to 0.25 ug/mL puromycin for 2 days. Antibiotic-resistant colonies were collected and expanded on Matrigel-coated plates in E8 medium. The mutation was confirmed using PCR product purification, subcloning into a TOPO TA cloning vector, and sequencing.

### Stem cell and organoid culture

Stem cell lines were maintained in six-well plates (NEST, 703003) coated with 1% Matrigel (Corning, 354277) in a 1:4 mixture of StemFlex and Essential 8 media (Thermo Fisher Scientific).The stem cells were cultured in an incubator, and the media changed every day until the cells reached 90% confluency. We followed our previously published protocol^43^ for generating SNR-derived organoids from control, patient, and isogenic control (H9) with MAST1-specific mutations including homozygous and heterozygous samples.

### Neuronal differentiation

After single neural rosettes were picked, neuroepithelial cells were expanded in neural differentiation (ND) medium, a 1:1 mixture of N2 medium (DMEM/F-12, 1% N2 supplement (cat#17502048), 1% MEM-NEAA, 2 µg/mL heparin and 1% penicillin/streptomycin) and B27 medium (Neurobasal-A (cat#10888022), 2% B27 supplement with vitamin A (cat#17504044), 1% GlutaMAX and 1% penicillin/streptomycin) in the absence of EGF and FGF2, with medium changed every 3-4 days. After 2-3 weeks, neurons were dissociated into single cells by incubation with Accutase (Stemcell, 07920) or papain (20 U/mL, Worthington) for 5-10 min at 37 °C. Trypsin inhibitor was then added to stop the enzyme reaction. Cells were passed through a 40 µm cell strainer (Corning, 431750) and plated on Matrigel-coated plates in a 1:1 mixture of ND medium. Depending on the experiment, iPSC-derived neurons were replated on Matrigel-coated 12-or 24-well plates, eight-chamber cell culture slides (CellTreat, 229168) or live-cell imaging dishes (FluoroDish, WPI, FD3510-100).

### Lentiviral preparation

The following lentivectors were designed and constructed. The WT-*MAST1* coding sequence (NM_014975.2_cds) (Gly-831-GGA to Gly-831-GGG) was cloned into the LentiCRISPR V2 (Addgene TLCV2 87360) vector between the AgeI/BamHI restriction sites, in front of a doxycycline-inducible promoter, tight TRE. The addition of doxycycline induced the production of *Mast1*-T2A-eGFP. In a similar fashion, *Mast1* mutant (c.3889 G>T, p.Asp1297Tyr) was cloned into the AgeI/BamHI restriction site. The negative control plasmid lacked the *Mast1* sequence, and only GFP was expressed under the doxycycline-inducible promoter. For morphology studies in which different cell lines were co-cultured, CamKIIα-GFP in a pLenti vector (Addgene plasmid # 96941) was used. GFP was substituted for RFP under the CamKIIα promoter. Lentiviruses were prepared as previously reported.

### Neuronal morphology analysis

Confocal z-stack 10X or 20X images of GFP-expressing neurons were acquired on Zeiss Airyscan and processed using Fiji/ImageJ (Version 2.16.0/1.54p; NIH) with the simple neurite tracer (SNT) plugin (v4.2.1). Maximum intensity projections were generated to obtain two-dimensional (2D) images, and neurites were manually traced using SNT to create skeletonized representations of neuronal morphology. Sholl analysis was performed using automated concentric circles centered on the neuronal soma with a fixed radial interval of 1 µm extending to distal neurite tips. Neurite intersections were quantified automatically. Each profile was generated from intersection counts, and parameters including the maximum number of intersections were extracted. Total neurite length per neuron was quantified from SNT-derived path length measurements by tracing all primary and secondary branches originating from the soma. This analysis was conducted by investigators blinded to the groups, and only GFP-positive neurons with visible soma were included.

### qRT-PCR

RNA was extracted from neurons and 1-month-old organoids with the Quick-RNA MicroPrep Kit (Cat#: R1050) according to the manufacturer’s protocol. Aliquots of 50 ng of total RNA were subjected to reverse transcription using the SuperScript™ III First-Strand Synthesis SuperMix (Cat#: 11752050). qRT-PCR assays were conducted using Applied Biosystems PowerUp SYBR Green Master Mix (Cat#: A25776) and the QuantStudio Real-Time PCR System (Applied Biosystems). Relative mRNA levels in each sample were normalized to the expression of h18S. Primers were designed using Primer Bank. Forward and reverse primers for *TUJ1* and *MAST1* were used. The sequences were: CGGTGGTGGAACCCTACAAC (*TUJ1*-forward), AGGTGGTGACTCCGCTCAT (*TUJ1*-reverse), CCGGCCAAACTGCACTCATC (*MAST1*-forward), and GATTCCTTCTCCTTGCTCGACA (*MAST1*-reverse).

### Bulk RNA Sequencing and Analysis

One-month-old SNR-derived organoids from Pat1 and the matched parental control Ctrl1 were pooled, with three organoids per sample and three samples per group. Total RNA was purified using the RNeasy Micro Kit (Qiagen) with DNase I digestion and assessed using the Bioanalyzer RNA Nano Assay (Agilent). Libraries were prepared using the NEBNext Ultra II Directional RNA Library Prep Kit with rRNA depletion and sequenced on an Illumina NovaSeq 6000 with 50-base paired-end reads.

The human GRCh38 genome and gene annotation files were downloaded from Ensembl release 100 and a reference database was created using STAR version 2.7.3a^63^. Optical duplicates were removed from the paired end FASTQ files using clumpify v38.34 and reads were trimmed of adapters using cutadapt 1.16^64^. The trimmed reads were aligned to the reference database using STAR in two pass mode to output a BAM file sorted by coordinates. Mapped reads were assigned to annotated genes using featureCounts version 1.6.3^65^.

Differentially expressed genes were identified using a 5% false discovery rate with DESeq2 version 1.28.1^66^. The regularized log values from DESeq2 were used in the principal components analysis and to create gene heatmaps. Developmental similarity was assessed by Spearman correlation with the BrainSpan dataset^67^. Cell-type proportions were estimated using cellular population mapping deconvolution analysis in the scBio package with a custom single cell dataset^68^.

### Single-cell RNA-sequencing

SNR-derived organoids (n=5/each group, iCtrl, Mut1 and Mut2) were rinsed with sterile 1X Dulbecco’s phosphate-buffered saline (DPBS) and chopped in small pieces using a surgical blade in the 30 cm petri dish. Chopped organoids were collected in 1.5 mL tubes in 1x DPBS and centrifuged at 1000 rpm for 2 min. Organoids samples were incubated in pre-warmed papain (Worthington biochemical) with 5% DNase (Sigma) for 20 min at 37C. After 6-10 times gentle pipetting again incubated 20 more min in incubator. After incubation trypsin inhibitor (Sigma) were used and then samples were centrifuged for 2 min at 1000 rpm. Added DPBS containing 0.04% BSA into the samples and were kept on ice. Samples were immediately transferred to the Genomics Core facility at the University of Utah.

### Library preparation and raw data processing for single-cell RNA-sequencing

Single-cell capture, lysis, and cDNA synthesis were performed at the University of Utah High Throughput Genomics Core using the Chromium system and Chromium Next GEM Single Cell 3′ Gene Expression v3.1 reagents (10x Genomics), according to the manufacturer’s protocol. Single-cell suspensions were loaded onto Chromium Single Cell 3′ chips with the appropriate master mix. Libraries were sequenced on an Illumina NovaSeq X Series instrument using a NovaSeq X Series 10B Reagent Kit with 2 × 150 bp paired-end reads (1,250 M read pairs per lane; Illumina). Sequencing reads were processed using the Cell Ranger pipeline (10x Genomics) and aligned to the human reference genome, GRCh38/hg38 (Homo sapiens, December 2013 annotation). Gene-barcode matrices were generated for each sample by counting unique molecular identifiers (UMIs).

### Data import and software environment

Single-cell RNA-sequencing data were analyzed using R version 4.5.2, Seurat version 5.5.1, and SeuratObject version 5.4.0. Seurat was configured to create version 3 assay objects. Cell Ranger-filtered feature–barcode matrices were imported separately for iCtrl, Mut1, and Mut2 using Read10X, and individual Seurat objects were generated using CreateSeuratObject with default settings. Cell barcodes were assigned sample-specific identifiers, and the objects were merged while retaining sample information. iCtrl was designated as Control, whereas Mut1 and Mut2 were grouped as MAST-mutant. The RNA assay was explicitly selected for downstream expression analyses. Unless otherwise specified, functions were run using their default settings.

### Quality control and cell filtering

Cells were assessed using detected gene counts, total unique molecular identifier (UMI) counts, and mitochondrial transcript percentage. Mitochondrial percentages were calculated using PercentageFeatureSet with the gene-name pattern "^MT-". Initial filtering retained cells with at least 1,500 detected genes, fewer than 350,000 UMIs, and less than 6% mitochondrial transcripts. These initial thresholds were intended to retain a broad population of cells for subsequent QC assessment. Cell counts and QC distributions were visualized by sample using violin plots, density plots, boxplots, and scatterplots. Relationships between detected genes and UMI counts were visualized separately by sample on log10 axes, with cells shaded by mitochondrial transcript fraction. Additional scatterplots displayed UMI counts against mitochondrial percentages and detected gene counts. Transcriptomic complexity was calculated as log10(detected genes) divided by log10(UMIs). Additional filtering retained cells with more than 2,000 UMIs, more than 2,000 detected genes, and a complexity score of at least 0.80. All three criteria were required for retention. Because these filters were applied to the initially filtered dataset, the upper UMI limit of 350,000 and mitochondrial limit of 6% remained in effect. The resulting dataset was used for normalization, integration, and doublet identification.

### Normalization and cell-cycle scoring

A random seed of 42 was set before preprocessing the filtered dataset. Gene expression was normalized using NormalizeData function with the LogNormalize method and a scale factor of 10,000. For each cell, gene counts were divided by the total RNA-assay count, multiplied by 10,000, and transformed using the natural logarithm of one plus the normalized value. S-phase and G2/M-phase scores were calculated using CellCycleScoring with the corresponding gene sets supplied in Seurat’s cc.genes object. These scores were retained as cell-level metadata and used as covariates during scaling.

### Variable-feature selection, scaling, and principal component analysis

The 2,000 most variable genes were selected using FindVariableFeatures with the variance-stabilizing transform method. Expression values were centered and scaled using ScaleData, with S-phase score, G2/M-phase score, and mitochondrial transcript percentage included as regression covariates. Principal component analysis was performed on the scaled variable genes using RunPCA with 50 components. Elbow plots were generated to assess the principal-component structure.

### Preliminary integration and doublet identification

Harmony version 2.0.5 was used to integrate the datasets by sample. Sample identity was supplied as the grouping variable to RunHarmony, with other settings left at their defaults. Preliminary UMAP visualization and nearest-neighbor graph construction used the first 15 Harmony dimensions. Preliminary graph-based clustering was performed using FindClusters with default settings. Putative doublets were identified using scDblFinder version 1.24.10. In the implemented workflow, the normalized RNA expression matrix was supplied as input, together with sample identifiers and the current cell identities through the clusters argument. Other settings were left at their defaults. Doublet scores and classifications were stored in the metadata, and cells classified as singlets were selected for subsequent processing.

### Final integration and graph-based clustering

Following singlet selection, normalization, cell-cycle scoring, variable-gene selection, scaling, principal component analysis, and Harmony integration were repeated using the settings described above. A shared nearest-neighbor graph was constructed using FindNeighbors with the first 15 Harmony dimensions. Final clustering was performed using FindClusters with the Louvain algorithm and a resolution of 0.175. Other neighbor-search and clustering settings were left at their defaults. The final analysis object contained 21,315 cells: 8,304 iCtrl cells, 6,670 Mut1 cells, and 6,341 Mut2 cells.

### UMAP visualization

UMAP embeddings were calculated using RunUMAP with the first 15 Harmony dimensions. Parameter combinations were explored using 10, 20, 30, or 40 neighbors; minimum distances of 0.05, 0.1, 0.3, 0.4, 0.5, or 0.6; and spread values of 1 or 3. The final visualization used 10 neighbors, a minimum distance of 0.1, and a spread of 1, with other settings left at their defaults. Cells were visualized by cluster, sample and genotype. Sample-and genotype-specific panels displayed subsets of the shared embedding rather than independently calculated embeddings.

### Cluster-marker identification and assessment of cluster identities

Cluster-associated genes were identified using FindAllMarkers with the Wilcoxon rank-sum test, comparing each cluster with the remaining cells. Testing used min.pct = 0.25 and logfc.threshold = 0.25, requiring detection in at least 25% of cells in either comparison group and a minimum absolute log2 fold change of 0.25. Other settings, including the function’s return threshold, were left at their defaults. Raw and Bonferroni-adjusted P values were retained. Returned genes were ranked within each cluster by average log2 fold change. The top 5, 10, and 100 markers were selected using this ranking and examined using heatmaps. Tied fold-change values could result in more than the specified number of markers being selected. No additional adjusted-P-value filter was applied when selecting these ranked lists. All clusters were manually assessed for consistency with known population-marker expression patterns using dot plots, feature plots, and violin plots. Marker expression was examined across clusters and separately by genotype or sample to support interpretation of cluster identities. The marker panels used for assessment are specified in the accompanying analysis code.

### Genotype differential-expression analysis

Gene expression was compared between Control and MAST-mutant cells using FindMarkers with the Wilcoxon rank-sum test. Comparisons were specified for the complete processed dataset and for individual clusters. Testing used min.pct = 0.1 and logfc.threshold = 0.25, requiring detection in at least 10% of cells in either group and a minimum absolute log2 fold change of 0.25. P values were adjusted using Seurat’s Bonferroni procedure based on the total number of genes in the analyzed assay. For comparisons of MAST-mutant cells against Control cells, positive fold changes indicate higher expression in MAST-mutant cells, whereas negative values indicated higher expression in Control cells. For reversed comparisons, the fold-change direction was reversed. The direction of each comparison was specified for the corresponding results. Cells were pooled by genotype without modeling sample-level replication. Multiple-testing correction was applied within each comparison, with no additional correction across separate cluster or subpopulation comparisons.

### Subpopulation selection and analysis

Marker-defined subpopulations were selected from the processed Seurat object independently of graph-based cluster assignments. Normalized expression values were extracted from the RNA assay for each selected marker or marker pair. Cells were included when log-normalized expression exceeded 1 for at least one selected marker. For paired-marker definitions, either marker could satisfy the expression threshold; co-expression was not required. Only markers present in the expression matrix were used, and subpopulations containing no qualifying cells were skipped. Exact marker definitions are provided in the accompanying figures or analysis code. Qualifying cells were selected regardless of genotype, and the original sample and genotype metadata were retained. Subpopulation assignments were nonexclusive, allowing individual cells to contribute to multiple subpopulations. Selected cells were analyzed using their existing normalized expression values without additional normalization, integration, or re-clustering.

### Differential expressions within subpopulations

Within each marker-defined subpopulation, MAST-mutant cells were compared with Control cells using FindMarkers with the Wilcoxon rank-sum test. Testing used min.pct = 0.1 and logfc.threshold = 0.25, requiring detection in at least 10% of cells in either comparison group and a minimum absolute log2 fold change of 0.25. Comparisons that could not be completed were skipped. Positive average log2 fold changes indicated higher expression in MAST-mutant cells, whereas negative values indicated higher expression in Control cells. Genes were classified as significant when the Bonferroni-adjusted P value was below 0.05 and the absolute average log2 fold change exceeded 0.25. Cells were pooled by genotype without modeling sample-level replication, and no additional multiple-testing correction was applied across separate subpopulation comparisons.

### Subpopulation average-expression scatterplots

Average RNA expression was calculated separately by genotype within each marker-defined subpopulation using AverageExpression with the RNA assay and normalized data layer. Log-normalized values were back-transformed before averaging across cells. The resulting values represented cell-weighted genotype averages rather than averages of sample-level means. Average-expression values were matched to all genes returned by the corresponding differential-expression analysis, rather than restricted to significant genes. Average Control expression was plotted on the x-axis and average MAST-mutant expression on the y-axis. Both axes used log10 scales with lower limits of 0.1; genes with average expression below this limit in either genotype were omitted from the displayed plot. A diagonal line indicated equal average expression. Significant genes were highlighted in red, and other returned genes were shown in gray. Significant genes were submitted for labeling using ggrepel, with label size 3, box padding 0.5, and a maximum overlap setting of 40. No additional statistical tests were performed for the scatterplots.

### Subpopulation composition plotting

The number of cells in each marker-positive subpopulation was calculated for Control and MAST-mutant groups. For each subpopulation, the cell count from each genotype was divided by the total cell count across both genotypes and multiplied by 100. The resulting percentages were displayed as stacked bar plots. Cells could contribute to more than one subpopulation if they met multiple marker-expression criteria. These comparisons were descriptive, and no statistical tests were performed.

### Gene Enrichment Analysis

Gene Ontology (GO) enrichment analysis was performed on differentially expressed genes identified within each marker-defined subpopulation using GO enrichment analysis tools^69^.

### Cluster expression similarity

Gene expression was aggregated within each cluster using the RNA assay. Cosine similarity was calculated between cluster-level expression profiles. The similarity matrix was ordered by hierarchical clustering using 1 minus the cosine similarity as the dissimilarity measure and displayed as a heatmap.

### Western blotting

Protein lysates from neurons collected after 2–3 weeks in differentiation conditions, 1-month-old organoids, and isolated mitochondrial and cytosolic fractions were prepared with 2X Laemmli Sample Buffer (Bio-Rad, cat.1610737). Protein lysates were loaded onto a NuPAGE 4–12% Bis-Tris gel (Invitrogen, cat. NP0336BOX) using a polyacrylamide gel electrophoresis (PAGE) chamber to separate protein samples. Separated proteins were transferred using the Trans-blot Turbo Transfer Pack (Bio-Rad, cat. 1704156) for 10 min at 4°C. Following transfer, membranes were blocked with 5% skim milk in PBS containing 0.1% Tween-20 (PBST) for 1 h while rocking at room temperature. Membranes were then incubated in solutions containing the following primary antibodies: anti-MAST1 (Novus, cat. NBP2-17228) at 1:1000, anti-HSP60 (R&D Systems, cat. MAB1800) at 0.5 μg/mL, anti-TUJ1 (Biolegend, cat. MMS-435P) at 0.1 µg/mL, anti-SIRT3 (Cell Signaling Technology, cat. 5490) at 1:1000, anti-TFAM (Cell Signaling Technology, cat. 8076) at 1:1000, anti-clv-Cas3 (Cell Signaling Technology, cat. 9661) at 1:1000, anti-TOMM20 (Cell Signaling Technology, cat. 42406) at 1:1000, anti-Oxphos complex (Abcam, cat. Ab110413) at 1:2000, and anti-β-actin (Proteintech, cat. 66009-1-lg) at 1:40,000 in 5% milk in PBST and slowly rocked at 4°C overnight. Membranes were washed three times for 10 min with PBST and then incubated with horseradish peroxidase (HRP)-conjugated secondary anti-rabbit (Cell Signaling Technology, cat. 7074) and anti-mouse (Cell Signaling Technology, cat. 7076) antibody at 1:3000 in 5% milk in PBST for 1 h and 30 min at room temperature. After three 10-min washes in PBST, the membranes were treated with chemiluminescent HRP substrate (EMD Millipore, cat. WBKLS0100) for protein visualization. Membranes were imaged with the ChemiDoc XRS + System and Image Lab Software (Bio-Rad, cat. 1708265), and the intensities of the detected protein bands were quantified using ImageJ (National Institutes of Health). Briefly, we used 10 µL protein lysates from each experimental group to obtain the β-actin bands, and the intensity values for these bands were used for normalization. Based on the normalization, we loaded the required amounts of protein lysates from different groups and finally measured the intensities of target protein bands.

### Immunohistochemistry and imaging of fixed cells and tissues

In preparation for immunohistochemical staining, 1-month-old SNR-derived organoids were fixed in 4% paraformaldehyde overnight at 4°C and washed with PBS. The organoids were then passed through a sequential sucrose solution gradient of 10%, 20%, and 30% separately, before being placed in cryosectioning molds, embedded in Optical Cutting Temperature (OCT) medium, flash frozen, and stored at –80°C. Frozen organoids were cut at 15–20 μm thickness using a Leica Cryostat machine and adhered to positively charged microscope slides. Cryosections were then processed for immunostaining by washing with 0.1M glycine in PBS and blocking with 0.3% Triton-X 100 with 3% BSA in PBS for 1–2 h at room temperature. Sections were then exposed to primary antibodies diluted in 0.3% Triton-X 100 with 3% BSA in PBS overnight in a cold room. The following primary antibodies were used: rabbit anti-PAX6 (Biolegend, PRB-278P 1:500), mouse anti-HOPX (Santa Cruz, sc-398703 1:200), rat anti-CTIP2 (Abcam, Ab18465 1:200), rabbit anti-GABA (Millipore, A2052 1:500), chicken anti-GFAP (Abcam, Ab4674 1:1000), rabbit anti-S100B (Invitrogen, PA5-78161 1:500), rabbit anti-MAST1 (Cusabio, CSB-PA897529LA01HU, 1:100), mouse anti-HSP60 (R&D Systems, MAB1800, 12 µg/mL), rabbit anti-TFAM (Cell Signaling Technology, #8076 1:100), chicken anti-TUJ1 (Novus, NB100-1612, 1:200), mouse, anti-Ki67 (BD Biosciences, 550609, 1:100), mouse anti-TOMM20 (Santa Cruz Biotechnology, sc-17764, 1:200), rabbit anti-caspase-3 (BD Pharmingen, 559565, 1:500), chicken anti-Map2 (Novus, NB300-213, 1:1000), mouse anti-SOX2 (Santa Cruz Biotechnology, sc-365823, 1:200), rabbit anti-TBR1 (Abcam, ab31940, 1:400), mouse anti-SATB2 (Abcam, ab51502, 1:500), rabbit anti-RFP (Evrogen, AB234, 1:1000), and chicken anti-GFP (Novus, NB100-1614, 1:500). The next day sections were washed with PBS for 15 min and exposed to secondary antibodies diluted in 0.3% Triton-X 100 with 3% BSA in PBS (1:1000). Sections were exposed to secondary antibodies coupled to Alexa Fluor fluorescent dyes (Invitrogen) and DAPI (1:1000) for 1–2 h at room temperature in darkness. After three 15-min PBS washes, slides were mounted with coverslips using Aqua Poly Mounting Media, dried at room temperature in darkness, and stored at –20°C for imaging. Fluorescence imaging was performed using a Zeiss 880 Airyscan confocal microscope at the University of Utah Cell Imaging Core Facility, and data processing and quantification were performed using Fiji and/or CellProfiler software.

### Mitochondrial respiration measurement

Stem cell-derived neurons and organoids were generated from H9 and MAST1-mutant engineered cell lines. Neurons were cultured for 2 weeks and treated with 10 µM AraC for 4–6 days. Then neurons were detached with papain and seeded at 50,000 cells/well in 200 μl N2/B27 media in 96-well plates (Agilent, Seahorse XF96) coated with a 1:100 dilution of Matrigel. Similarly, organoids were transferred to 96-well Seahorse plates coated with Matrigel. Neurons and organoids were incubated for 48 h at 37°C in a 5% CO_2_ incubator. The oxygen consumption rate (OCR) was determined using the Seahorse XF96 Extracellular Flux Analyzer with sequential addition of oligomycin (1 μM), carbonyl cyanide 4-trifluoromethoxy phenylhydrazone (FCCP) (3 μM), rotenone (0.5 μM), and antimycin (0.5 μM) to measure cellular respiration. OCR measurements were normalized to gDNA.

### Fluorescence-activated cell sorting (FACS)

For detection of mito-tracker green and red by FACS, 1-month-old iCtrl and MAST1-mutant organoids were washed with 1X PBS and treated with 500 µl accutase and 25 µl DNAse. After gentle pipetting 12–5 times, the organoids were incubated at 37°C for 5 min and then again pipetted gently 20 times. If dissociation into single cells was still not achieved, they were returned to the incubator for an additional 2 min. Finally, 500 µl trypsin inhibitor was added, and samples were centrifuged at 2000 rpm for 2 min. After washing with FACS buffer, centrifugation was repeated. Then the cells were filtered with a cell strainer into single cells and separated for preparation of unstained control and stained samples. Mito tracker green and Mito tracker red were added, and the cells were incubated at 37°C for 20 min. After two washes with FACS buffer, samples were centrifuged, and the supernatant was aspirated before addition of 250 µl FACS buffer. The final samples were placed on ice and covered with foil for immediate transfer to the core lab flow cytometry facility.

For ROS measurement by FACS, 2–3-week-old MAST1-mutant neurons and patient iPSC-derived neurons were dissociated following standard procedures and treated with 5–10 µM of H2DCFDA (Invitrogen, cat. no. D399) for 20 min at 37°C in an incubator with 5% CO_2_. After collection by centrifugation and washing with 1X PBS containing 0.04% BSA, the samples were placed on ice and transferred to the core FACS facility.

### Transmission electron microscopy (TEM)

For viewing by TEM, 1-month-old organoids were collected and immersed in primary fixative solutions containing 1% glutaraledehyde, 2.5% paraformaldehyde, 100 mM cacodylate buffer, pH 7.4, 6 mM CaCl_2_ and 4.8% sucrose overnight at 4°C. The next day the samples were washed for 15 min with cacodylate buffer before secondary fixation using 2% osmium tetroxide at room temperature for 1 h. After two washes with cacodylate buffer and one 15-min wash with dH_2_O, samples were stained with saturated uranyl acetate for 1 h at room temperature and dehydrated with a graded ethanol series (30%, 50%, 70%, 95% two times, and 100% three times for 15 min each) followed by absolute acetone (10 min, three times). Organoids samples were embedded by immersion in 30% Epon epoxy resin solution for 5 h followed by 70% solution overnight and 100% resin for 8 h with three changes before immersion in fresh resin for 48 h at 60°C. Fixed and embedded organoid samples were ultracut using a Leica UC 6 ultratome at 70-nm thickness and mounted on 200 mesh copper grids for staining with saturated uranyl acetate for 20 min and lead citrate for 10 min. Finally, organoid sections were examined and imaged at an accelerating voltage of 120 kV on a JEOL-1400 plus transmission electron microscope (JEOL, Japan) equipped with a CCD Gatan camera. The average mitochondrial length and number of mitochondria per ROI were measured using imageJ.

### Mitochondrial isolation

One-month-old organoids were collected in 6-cm Petri dishes and rinsed with sterile 1X DPBS before being cut into small pieces with a surgery blade. One portion of the small pieces was collected for preparation of total cell lysates using Laemmli sample buffer (Bio-Rad, #1610737). Then the remaining small pieces were used for mitochondrial isolation using the Mitochondria Isolation Kit from Thermo Scientific (#89874) according to the manufacturer’s instructions. Based on the experimental plan, one mitochondrial fraction samples was treated with 100 µg/mL proteinase K (Bio Labs, #P8102S) at 50°C for 30 min before washing with mitochondria isolation reagent C (500 µL), centrifugation at 3000×g for 5 min, and lysis with Laemmli sample buffer. The other mitochondrial fraction was treated with Laemmli sample buffer without proteinase K treatment. All samples were stored frozen at –20°C for use in further analyses.

### Live cell imaging

Neurons were transduced with 10 µL per well of lentivirus encoding CaMKII::tRFP_T2A_mito-Emerald, added to 6-well plates after removing most of the culture medium to leave a minimal volume and prevent drying, and incubated for 24 h at 37 °C and 5% CO₂. Medium was then replaced with fresh N2/B27 medium and changed every 3 days. One week after infection, neurons were dissociated with papain, replated onto Matrigel-coated live-cell imaging dishes at 6 × 10⁴ cells per well, and cultured for a further 5-7 days before imaging. Neurons were maintained at 37°C in 5% CO2, and time-lapse imaging was performed using a confocal microscope (Nikon Ring TIRF or Spinning Disk Confocal) and Keyence microscope. For dual-channel time-lapse imaging, images of both neurons (CamKII::tRFP) and mitochondria (mito-Emerald) were taken every 1 sec with green and red channels for a total duration of 5 min in the spinning disk microscope. Similarly, dual-channel time-lapse images of both neurons (red) and mitochondria (green) were taken every 30 sec for a total duration of 1 h in the Keyence microscope. Kymographs were generated with extra plug-ins for ImageJ. Live imaging movies were quantitatively analyzed using Fiji/ImageJ (v1.54p; 17 February 2025) by generating kymographs along defined linear axonal segments, using the KymographClear 2.0 plugin. Trajectories of mitochondrial trafficking were traced on kymographs manually and then segmented into continuous runs. For each continuous run, the run length (distance traveled along the track) and run time (cumulative time in directed motion) were measured, and the run velocity was calculated as run length divided by run time. Objects were classified as motile using a velocity threshold, enabling per-object summaries: time mobile versus stationary, mean speed while moving, mean run length, and the fraction of objects moving. Event frequencies were reported as counts normalized to both track length and acquisition time. Statistical comparisons were performed using standard two-tailed t-tests in GraphPad Prism (v11.0.0).

### Axon tract formation assay

Tissue chips were kindly provided by Dr. Yoshiho Ikeuchi (The University of Tokyo). Prior to use, tissue chips were submerged in 70% ethanol overnight. Then the chips were placed on tissue slides and pressed down gently. Tissue chips were allowed to completely dry and attach to the slides overnight. The tissue device channel was coated with 5% Matrigel for 2 h at 37°C. After 2 h, the coating solution was removed with P1000 and P200 pipettes through the channel without air bubbles. Two iCtrl organoids (1.5 months old) from the same differentiation batch were placed in opposing wells connected by a Matrigel-coated microchannel (n=16 organoids). Similarly, two age-matched MAST1-mutant organoids from the same batch were placed in opposing wells connected by a Matrigel-coated microchannel (n=16 organoids). Organoids are fed on day 1 and then twice a week with N2+B27 media (1% Matrigel). The medium volume was 200 µl per well. After 3 months in culture, axon tracts extending through the microchannels were physically separated from the cell body regions using sterile surgical blades. Axonal and somatic compartments were collected separately into microcentrifuge tubes and processed for protein lysis and immunoblot analysis.

### Pharmacology treatment of neurons and organoids

Control, patient, isogenic control (iCtrl) and MAST1-mut neural rosettes were manually picked and transferred into 96-well plates. Pharmacological treatment started 5-7 days after picking. Organoids and neurons were treated with creatine (4 mM) or N-acetylcysteine (2 mM) to assess rescue of organoid growth and neuronal morphology. Organoids were treated for 50 days and neurons for 3-4 weeks, with culture medium (N2/B27) replaced every other day throughout. After 50 days of treatment, organoid samples were prepared for immunoblotting. Neurons were cultured on eight-well chamber slides and, after 3-4 weeks of treatment, were fixed with 4% paraformaldehyde (PFA) and immunostained for MAP2 with DAPI. Confocal images were then acquired for neuronal morphology analysis.

### Immunofluorescence image analysis

Image analysis was performed blinded to experimental conditions. Image identifiers were randomized using a Python script, and condition labels were revealed after measurements were finalized. Images were analyzed in Fiji/ImageJ (v1.54p). Z-stacks comprising four optical sections were converted to maximum-intensity projections, normalized for brightness and contrast, and processed using a median filter (radius, 4). Quantifications were performed within square regions of interest (ROIs) of defined dimensions, with placement guided by DAPI and the relevant marker channels. Marker-positive objects were segmented using global Otsu thresholding and counted using Analyze Particles.

### DAPI-based object segmentation and marker quantification

A custom Fiji macro processed DAPI-derived binary masks within selected ROIs. Masks were restricted to ROI boundaries and separated by watershed, and objects were detected using Analyze Particles with size and circularity filtering. Each detected object ROI was applied to the corresponding marker channels to measure mean fluorescence intensity. Intensities were pooled across datasets separately for each marker, and marker positivity was defined as intensity ≥ the pooled median + 2 standard deviations. Marker-positive object counts were summarized per ROI.

### Segmented object-area measurements

ROIs were placed in progenitor zones using DAPI and SOX2 labeling. Marker-positive objects were segmented using global Otsu thresholding, and projected object area (μm²) was measured using Analyze Particles. SOX2 measurements were restricted to objects with fluorescence intensity in the top decile (≥90th percentile), a criterion selected by comparison with manual assessment.

### Image quantification using Cell Quant Studio

Images were analyzed using Cell Quant Studio (v0.1), a custom Fiji/ImageJ workflow implemented through a Python interface and ImageJ macros. Four-channel images were processed into channel-specific, two-dimensional measurement images using configurable projection or slice-selection and filtering settings. A binary segmentation mask was generated from DAPI labeled channel using intensity thresholding and watershed separation. Within user-defined ROIs, segmented objects were detected using Analyze Particles with specified area and circularity limits. Each object ROI was applied to all four measurement channels to extract mean fluorescence intensity and integrated density. Object area, perimeter, circularity and Feret diameter were also measured. Object-level measurements and ROI-level counts and total segmented areas were exported as CSV files for downstream analysis. Source code is available at https://github.com/KiViKatt/ImageJ-SingleCellQuantificationPlugin.

### Experiments with Drosophila melanogaster

The following fly lines were used: *EGFP* RNAi (*P{VALIUM22-EGFP.shRNA.1}attP40*), *dop* RNAis (y[1] sc[*] v[1] sev[21]; P{y[+t7.7] v[+t1.8]=TRiP.HMS05876}attP40) (y[1] sc[*] v[1] sev[21]; P{y[+t7.7] v[+t1.8]=TRiP.HMC05132}attP40) (y[1] v[1]; P{y[+t7.7] v[+t1.8]=TRiP.JF02778}attP2) (y[1] v[1]; P{y[+t7.7] v[+t1.8]=TRiP.HMC02943}attP40), *inscuteable-GAL4* (*P{w[+mW.hs]=GawB}insc[Mz1407]), neuronal Synaptobrevin-GAL4* (*P{y[+t7.7] w[+mC]=nSyb-GAL4.P}attP2*), *dop[1] red[1] e[1]/TM6B, Tb[1*] (PMID: 24803657), *(P{w[+mW.hs]=GAL4-da.G32}UH1), y[1] w[*] PBac{y[+mDint2] w[+mC]=UAS-human MAST1.HA}VK00037, y[1] w[*] PBac{y[+mDint2] w[+mC]=UAS-human MAST1pmut.HA}VK00037, y[1] w[*] PBac{y[+mDint2] w[+mC]=UAS-human MAST1mitomut.HA}VK00037*. All flies were maintained at 25°C and grown on Archon glucose formula food in medium, wide plastic vials. RNAi crosses were grown at 29°C on Archon glucose formula food with bromophenol blue. Brain volume measurements were taken on wandering third-instar larval brains. The wandering third-instar larval stage was identified morphologically by extruding spiracles and gut clearance. Brains were analyzed using the Imaris software. A new surface was generated on the Deadpan channel, and the brain was traced every 5^th^ z stack. We then created a surface to compile all traces, and volume was logged from the results section. Volumes were analyzed using GraphPad Prism software using a multiple comparison ANOVA which compared the mean from each *dop* RNAi group to the mean of the *EGFP* RNAi control.

### Quantification and statistical analysis

All experimental materials and cell lines including human iPSCs and engineered lines used in the different experiments are included in the Supplementary Table. Statistical tests and sample sizes are indicated in the figure legends. All neurons and organoids were randomly selected for use in experiments. Excel and GraphPad Prism 10 were used for data analysis and statistical testing.

## Acknowledgements

We thank D.Viskochil for assistance with patient assessments; T. Tvrdik for assistance with patient recruitments; M. Deshotel for assistance with sequencing and variant validation; N. Romero, C. Russell, C. Armstrong, and K. Burger for assistance with stem cell and organoid cultures; S. Flat for assistance with image quantification; C. Goh for assistance with imaging experiments. The study was supported by the NINDS (R01NS123849 and R01NS149744) and Utah Genome Project grants (to A.S.) and NIH Genomic Medicine Training Grant (T32HG008962) (to H.M.A.U.).

## Notes

### Competing Interest Statement

The authors have declared no competing interest.

## References

1. Romero-Morales, A. I. & Gama, V. Revealing the Impact of Mitochondrial Fitness During Early Neural Development Using Human Brain Organoids. Front. Mol. Neurosci. 15, (2022).

2. Iwata, R., Casimir, P. & Vanderhaeghen, P. Mitochondrial dynamics in postmitotic cells regulate neurogenesis. Science (80-. ). 369, 858–862 (2020).

3. Pekkurnaz, G. & Wang, X. Mitochondrial heterogeneity and homeostasis through the lens of a neuron. Nat. Metab. 4, 802–812 (2022).

4. Manji, H. et al. Impaired mitochondrial function in psychiatric disorders. Nat. Rev. Neurosci. 13, 293–307 (2012).

5. Schon, E. A. & Przedborski, S. Mitochondria: The Next (Neurode)Generation. Neuron 70, 1033–1053 (2011).

6. Misgeld, T. & Schwarz, T. L. Mitostasis in Neurons: Maintaining Mitochondria in an Extended Cellular Architecture. Neuron 96, 651–666 (2017).

7. Pilling, A. D., Horiuchi, D., Lively, C. M. & Saxton, W. M. Kinesin-1 and Dynein Are the Primary Motors for Fast Transport of Mitochondria in Drosophila Motor Axons. Mol. Biol. Cell 17, 2057–2068 (2006).

8. van Spronsen, M. et al. TRAK/Milton Motor-Adaptor Proteins Steer Mitochondrial Trafficking to Axons and Dendrites. Neuron 77, 485–502 (2013).

9. MacAskill, A. F. et al. Miro1 Is a Calcium Sensor for Glutamate Receptor-Dependent Localization of Mitochondria at Synapses. Neuron 61, 541–555 (2009).

10. Kang, J. S. et al. Docking of Axonal Mitochondria by Syntaphilin Controls Their Mobility and Affects Short-Term Facilitation. Cell 132, 137–148 (2008).

11. Bapat, O. et al. VAP spatially stabilizes dendritic mitochondria to locally support synaptic plasticity. Nat. Commun. 15, 1–18 (2024).

12. Tröger, J. et al. Mitochondria are positioned at dendritic branch induction sites, a process requiring rhotekin2 and syndapin I. Nat. Commun. 16, (2025).

13. Pekkurnaz, G., Trinidad, J. C., Wang, X., Kong, D. & Schwarz, T. L. Glucose regulates mitochondrial motility via Milton modification by O-GlcNAc transferase. Cell 158, 54–68 (2014).

14. Courchet, J. et al. Terminal axon branching is regulated by the LKB1-NUAK1 kinase pathway via presynaptic mitochondrial capture. Cell 153, 1510 (2013).

15. Rangaraju, V., Lauterbach, M. & Schuman, E. M. Spatially Stable Mitochondrial Compartments Fuel Local Translation during Plasticity. Cell 176, 73–84.e15 (2019).

16. López-Doménech, G. et al. Loss of Dendritic Complexity Precedes Neurodegeneration in a Mouse Model with Disrupted Mitochondrial Distribution in Mature Dendrites. Cell Rep. 17, 317–327 (2016).

17. Barel, O. et al. Deleterious variants in TRAK1 disrupt mitochondrial movement and cause fatal encephalopathy. Brain 140, 568–581 (2017).

18. Liu, Y.-T. et al. Extended phenotypic spectrum of KIF5A mutations. Neurology 83, 612–619 (2014).

19. Poirier, K., Lebrun, N., Broix, L. & et. al. Mutations in TUBG1, DYNC1H1, KIF5C and KIF2A cause malformations of cortical development and microcephaly. Nat. Genet. 45, 639–47 (2013).

20. Nicolas, A. et al. Genome-wide Analyses Identify KIF5A as a Novel ALS Gene. Neuron 97, 1268–1283.e6 (2018).

21. Lindhout, F. W., Krienen, F. M., Pollard, K. S. & Lancaster, M. A. A molecular and cellular perspective on human brain evolution and tempo. Nature 630, 596–608 (2024).

22. Mohan, H. et al. Dendritic and axonal architecture of individual pyramidal neurons across layers of adult human neocortex. Cereb. Cortex 25, 4839–4853 (2015).

23. Iwata, R. et al. Mitochondria metabolism sets the species-specific tempo of neuronal development. Science (80-. ). 379, (2023).

24. Tripathy, R. et al. Mutations in MAST1 Cause Mega-Corpus-Callosum Syndrome with Cerebellar Hypoplasia and Cortical Malformations. Neuron 100, 1354–1368.e5 (2018).

25. Lumeng, C. et al. Interactions between β2-syntrophin and a family of microtubule-associated serine/threonine kinases. Nat. Neurosci. 2, 611–617 (1999).

26. Lemke, M. C. et al. MAST Kinases’ Function and Regulation: Insights from Structural Modeling and Disease Mutations. Biomedicines 13, (2025).

27. McMichael, G. et al. Whole-exome sequencing points to considerable genetic heterogeneity of cerebral palsy. Mol. Psychiatry 20, 176–182 (2015).

28. Bowling, K. M. et al. Genomic diagnosis for children with intellectual disability and/or developmental delay. Genome Med. 9, 1–11 (2017).

29. Gilissen, C. et al. Genome sequencing identifies major causes of severe intellectual disability. Nature 511, 344–347 (2014).

30. Kaplanis, J. et al. Evidence for 28 genetic disorders discovered by combining healthcare and research data. Nature 586, 757–762 (2020).

31. Dolan, M. et al. A novel microdeletion/microduplication syndrome of 19p13.13. Genet. Med. 12, 503–511 (2010).

32. Ben-Mahmoud, A., Al-Shamsi, A. M., Ali, B. R. & Al-Gazali, L. Evaluating the Role of MAST1 as an Intellectual Disability Disease Gene: Identification of a Novel De Novo Variant in a Patient with Developmental Disabilities. J. Mol. Neurosci. 70, 320–327 (2020).

33. Hecher, L. et al. The Clinical Picture of a Bilateral Perisylvian Syndrome as the Initial Symptom of Mega-Corpus-Callosum Syndrome due to a MAST1-Gene Mutation. Neuropediatrics 51, 435–439 (2020).

34. Rodríguez-García, M. E. et al. MAST1 variant causes mega-corpus-callosum syndrome with cortical malformations but without cerebellar hypoplasia. Am. J. Med. Genet. Part A 182, 1483–1490 (2020).

35. Antonioli, S. et al. A complex of MAST1 and 14-3-3η regulates Tau phosphorylation in the developing cortex. bioRxiv 2025.07.09.663707 (2025) doi:10.1073/pnas.2617534123/-/DCSupplemental.Published.

36. An, S. W. A. et al. KIN-4/MAST kinase promotes PTEN-mediated longevity of Caenorhabditis elegans via binding through a PDZ domain. Aging Cell 18, (2019).

37. Hain, D. et al. The Drosophila MAST kinase drop out is required to initiate membrane compartmentalisation during cellularisation and regulates dynein-based transport. Dev. 141, 2119–2130 (2014).

38. Jin, L. et al. MAST1 Drives Cisplatin Resistance in Human Cancers by Rewiring cRaf-Independent MEK Activation. Cancer Cell 34, 315–330.e7 (2018).

39. Buljan, M. et al. Kinase Interaction Network Expands Functional and Disease Roles of Human Kinases. Mol. Cell 79, 504–520.e9 (2020).

40. Strober, B. J. et al. The GTEx Consortium atlas of genetic regulatory effects across human tissues. Science (80-. ). 18, 1318–1330 (2020).

41. Li, M. et al. Integrative functional genomic analysis of human brain development and neuropsychiatric risks. Science (80-. ). 362, (2018).

42. Link, N. et al. Mutations in ANKLE2, a ZIKA Virus Target, Disrupt an Asymmetric Cell Division Pathway in Drosophila Neuroblasts to Cause Microcephaly. Dev. Cell 51, 713–729.e6 (2019).

43. Ullah, H., Huang, Q., Chiola, S., Wang, Y. & Shcheglovitov, A. Generating and Characterizing Human Telencephalic Brain Organoids from Stem Cell-Derived Single Neural Rosettes. Nat. Protoc. (accepted), (2025).

44. Wang, Y. et al. Modeling human telencephalic development and autism-associated SHANK3 deficiency using organoids generated from single neural rosettes. Nat. Commun. (2022) doi:10.1038/s41467-022-33364-z.

45. Alcamo, E. A. et al. Satb2 Regulates Callosal Projection Neuron Identity in the Developing Cerebral Cortex. Neuron 57, 364–377 (2008).

46. Martins-Costa, C. et al. ARID1B controls transcriptional programs of axon projection in an organoid model of the human corpus callosum. Cell Stem Cell 31, 866–885.e14 (2024).

47. Osaki, T. et al. Three-dimensional motor nerve organoid generation. J. Vis. Exp. 2020, 1–18 (2020).

48. Nowakowski, T. J. et al. Spatiotemporal gene expression trajectories reveal developmental hierarchies of the human cortex. Science (80-. ). 358, 1318–1323 (2017).

49. Yang, G. et al. Neurite outgrowth deficits caused by rare PLXNB1 mutation in pediatric bipolar disorder. Mol. Psychiatry (2023) doi:10.1038/s41380-023-02035-w.

50. Shcheglovitov, A. et al. SHANK3 and IGF1 restore synaptic deficits in neurons from 22q13 deletion syndrome patients. Nature 503, 267–271 (2013).

51. Redza-Dutordoir, M. & Averill-Bates, D. A. Activation of apoptosis signalling pathways by reactive oxygen species. Biochim. Biophys. Acta - Mol. Cell Res. 1863, 2977–2992 (2016).

52. Fukumitsu, K. et al. Synergistic action of dendritic mitochondria and creatine kinase maintains ATP homeostasis and actin dynamics in growing neuronal dendrites. J. Neurosci. 35, 5707–5723 (2015).

53. Samuni, Y., Goldstein, S., Dean, O. M. & Berk, M. The chemistry and biological activities of N-acetylcysteine. Biochim. Biophys. Acta 1830, 4117–4129 (2013).

54. Sheng, Z.-H. & Cai, Q. Mitochondrial transport in neurons: impact on synaptic homeostasis and neurodegeneration. Nat. Rev. Neurosci. 13, 77–93 (2012).

55. Beaulieu-Laroche, L. et al. Enhanced Dendritic Compartmentalization in Human Cortical Neurons. Cell 175, 643–651.e14 (2018).

56. Gidon, A. et al. Dendritic action potentials and computation in human layer 2/3 cortical neurons. Science 367, 83–87 (2020).

57. Spalding, K. L., Bhardwaj, R. D., Buchholz, B. A., Druid, H. & Frisén, J. Retrospective birth dating of cells in humans. Cell 122, 133–143 (2005).

58. Wyss, M. & Kaddurah-Daouk, R. Creatine and creatinine metabolism. Physiol. Rev. 80, 1107–1213 (2000).

59. Han, Q. et al. Restoring Cellular Energetics Promotes Axonal Regeneration and Functional Recovery after Spinal Cord Injury. Cell Metab. 31, 623–641.e8 (2020).

60. Henderson, J. T., Javaheri, M., Kopko, S. & Roder, J. C. Reduction of lower motor neuron degeneration in wobbler mice by N-acetyl-L-cysteine. J. Neurosci. 16, 7574–7582 (1996).

61. Qian, H.-R. & Yang, Y. Neuron differentiation and neuritogenesis stimulated by N-acetylcysteine (NAC). Acta Pharmacol. Sin. 30, 907–912 (2009).

62. Andres, R. H. et al. Effects of creatine treatment on the survival of dopaminergic neurons in cultured fetal ventral mesencephalic tissue. Neuroscience 133, 701–713 (2005).

63. Dobin, A. et al. STAR: ultrafast universal RNA-seq aligner. Bioinformatics 29, 15–21 (2013).

64. Kechin, A., Boyarskikh, U., Kel, A. & Filipenko, M. cutPrimers: A New Tool for Accurate Cutting of Primers from Reads of Targeted Next Generation Sequencing. J. Comput. Biol. a J. Comput. Mol. cell Biol. 24, 1138–1143 (2017).

65. Liao, Y., Smyth, G. K. & Shi, W. featureCounts: an efficient general purpose program for assigning sequence reads to genomic features. Bioinformatics 30, 923–930 (2014).

66. Love, M. I., Huber, W. & Anders, S. Moderated estimation of fold change and dispersion for RNA-seq data with DESeq2. Genome Biol. 15, 1–21 (2014).

67. Miller, J. A. et al. Transcriptional landscape of the prenatal human brain. Nature 508, 199–206 (2014).

68. Frishberg, A. et al. Cell composition analysis of bulk genomics using single cell data. Nat. Methods 16, 327–332 (2019).

69. Mi, H. et al. Protocol Update for large-scale genome and gene function analysis with the PANTHER classification system (v.14.0). Nat. Protoc. 14, 703–721 (2019).

