## Supplemental Data 1 for "MAST1 is neuron-specific regulator of mitochondrial trafficking and abundance in human brain tissue"

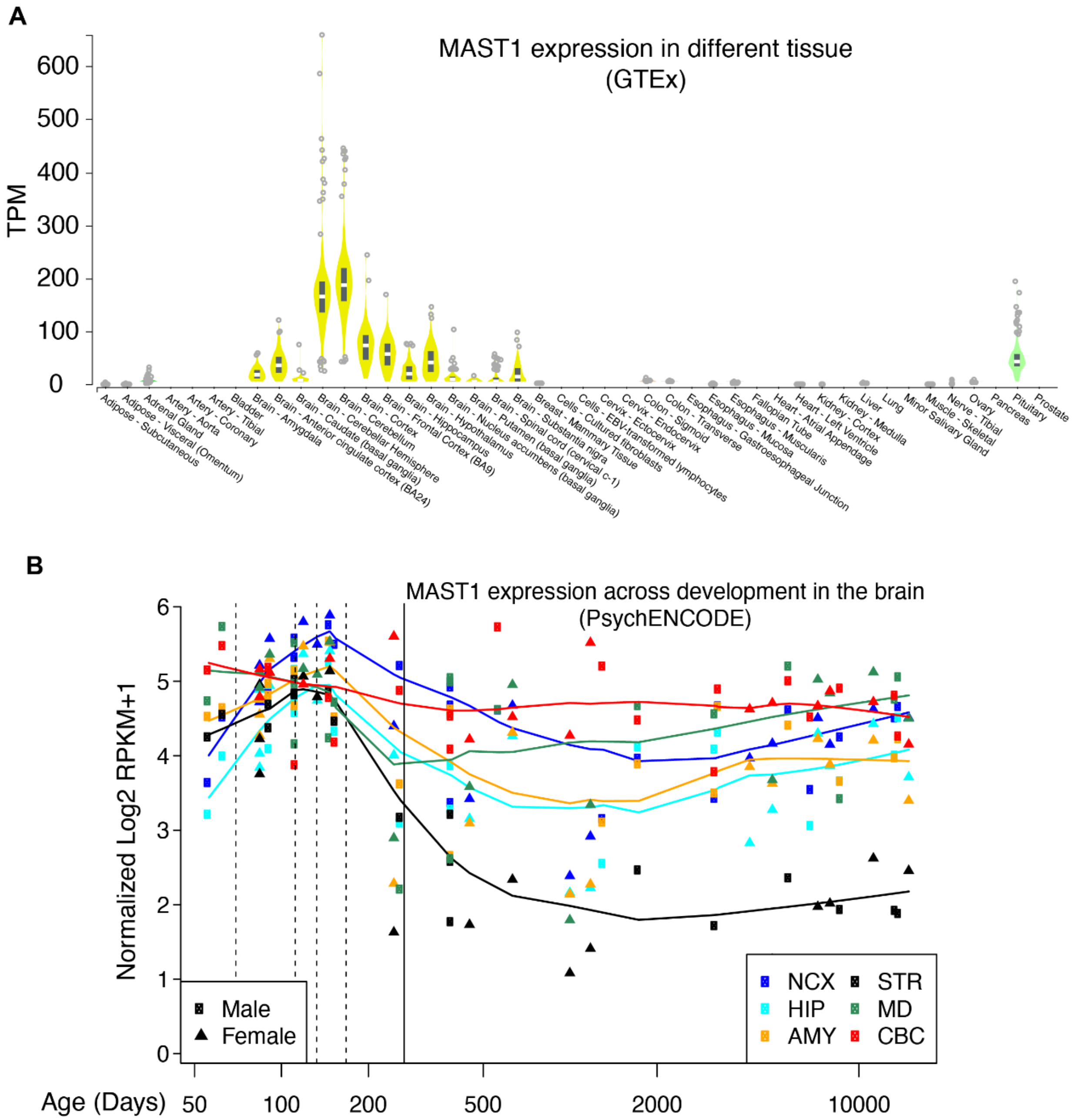


**Supplementary Figure 1. *MAST1* expression in adult human tissues and throughout human brain development. (A)** Violin plots of *MAST1* expression (transcripts per million, TPM) across human tissues from the Genotype-Tissue Expression (GTEx) project. *MAST1* is expressed predominantly in the brain, with the highest levels in the cerebellar hemisphere and cerebellum. Boxes indicate the median and interquartile range; dots represent individual samples. **(B)** *MAST1* expression (normalized log2 RPKM+1) across human brain development (PsychENCODE). Data are shown for neocortex (NCX), hippocampus (HIP), amygdala (AMY), striatum (STR), mediodorsal nucleus of the thalamus (MD) and cerebellar cortex (CBC). Squares denote male and triangles female samples; lines are smoothed fits per region. Age is plotted in days (log scale); the solid vertical line marks birth and dashed lines mark successive prenatal developmental windows. *MAST1* expression peaks prenatally and declines postnatally in most regions, most markedly in the striatum.

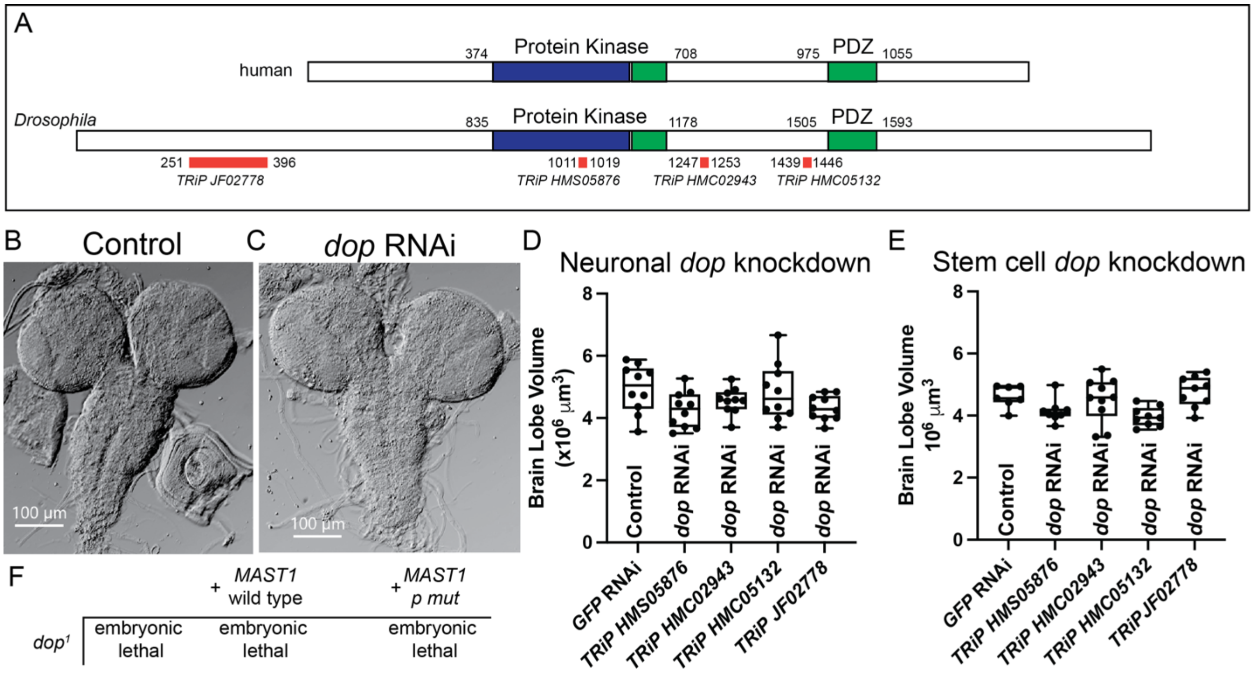


**Supplementary Figure 2. Loss of the *Drosophila* ortholog of *MAST1*, *dropout* (*dop*), does not significantly affect brain size. (A)** The protein structure for human MAST1 and the *Drosophila* ortholog, Dop, with conserved domains marked and RNAi targeting amplicons marked in orange. **(B-C)** Brightfield images of third-instar larval brains from a control RNAi knockdown of *EGFP* (B) and *dop* (C) in neurons. **(D)** Brain lobe volume measurements from animals expressing RNAi targeting *EGFP* (control) or *dop* in larval neurons using *n-Syb-GAL4*. Brain lobe volume did not differ significantly with neuronal knockdown of *dop* versus *EGFP*. **(E)** Brain lobe volume measurements from animals expressing RNAi targeting *EGFP* (control) or *dop* in larval neural stem cells using *insc-GAL4*. Brain lobe volume did not differ significantly with neural stem cell knockdown of *dop* versus *EGFP* except for line TRiP HMC05132 (*P=0.0383). One-Way ANOVA compared the mean of each dop RNAi line to the mean of the control; neurons: F=2.136, df=49, P=0.0918, neural stem cell: F=4.031, df=42, P=0.008. (F) WT and mutant Pro1177Ser mutant human *MAST1* were expressed in *dop*$^{1}$ mutants to test for rescue and conservation of function. Human transgene expression with the ubiquitous driver *da-GAL4* could not rescue embryonic lethality of *dop* loss.

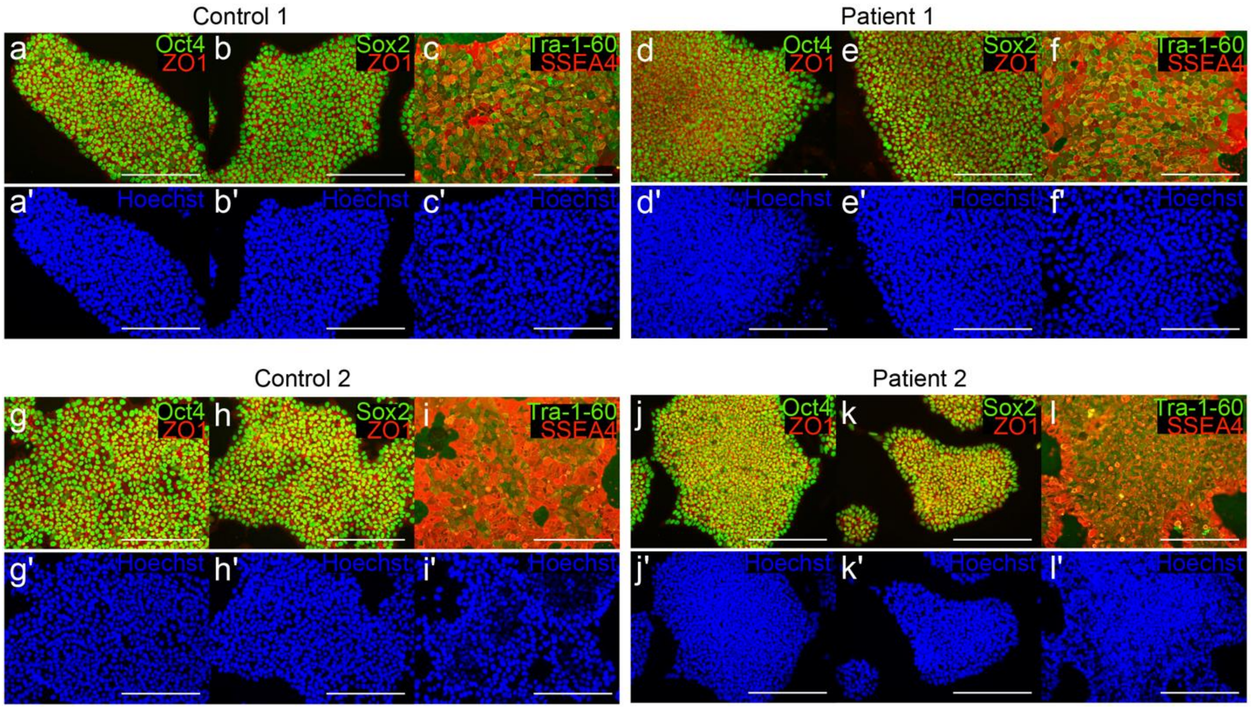


**Supplementary Figure 3. Characterization of iPSCs from controls and patients.** Immunofluorescence staining of pluripotency markers in iPSC lines from control 1 **(a-c')**, patient 1 **(d-f')**, control 2 **(g-i')** and patient 2 **(j-l')**. Colonies were stained for OCT4 (green) with the tight junction protein ZO1 (red) (a, d, g, j); SOX2 (green) with ZO1 (red) (b, e, h, k); and TRA-1-60 (green) with SSEA4 (red) (c, f, i, l). Nuclei were counterstained with Hoechst (blue; a'-l'). All lines uniformly expressed pluripotency markers. Scale bars, 200 µm.

### Generating CRISPR-engineered lines

#### A. Oligonucleotides used in this study

| Name | Type | Sequence (5'-3') |
|---|---|---|
| MAST1-KI-sgRNA2 | CRISPR sgRNA | GCTCGCCATGGAAGTCCTTG |
| Donor (Forward) | HDR donor (ssODN) | AGGGCGCGCTGCGCAAACACAGCCTCGAGGTGGG CCACCCGGATTTCCGCAAG**T**ACTTCCATGGCGAGCT GGCGCTGCATAGCCTTGCCGAGTCCGAC |
| PCR primer (forward) | Genotyping PCR | CCGGCCAAACTGCACTCATC |
| PCR primer (revrse) | Genotyping PCR | GATTCCTTCTCCTTGCTCGACA |

#### B. Human MAST1 locus (graphics made based on NM_014975.3)

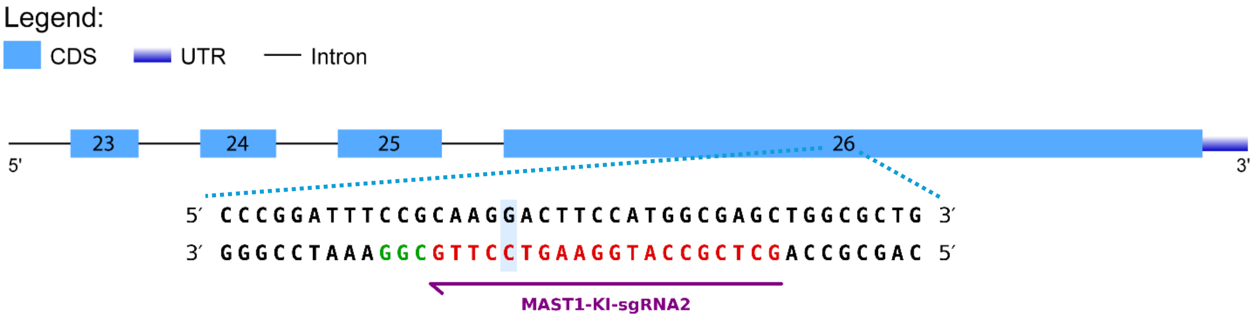


#### C. Representative sequencing chromatograms

**iCtrl (H9)**

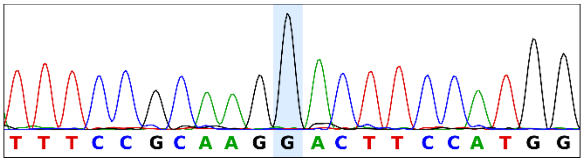


**Mut1 (Het21)**

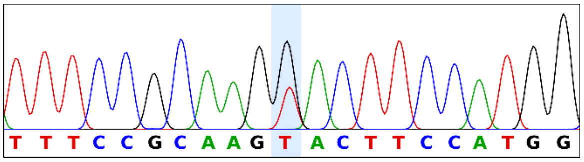


**Mut2 (Het73)**

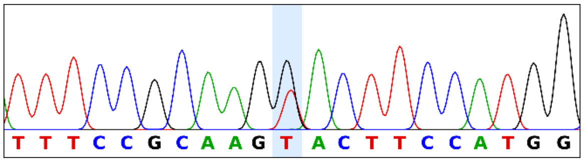


**Homozygous**

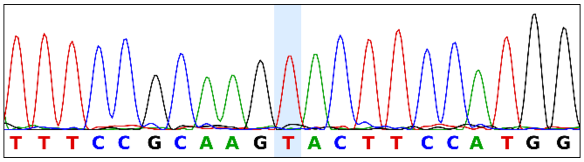


**Supplementary Figure 4. Generation of CRISPR/Cas9-engineered MAST1 mutant lines. (A)** Oligonucleotides used in this study: the CRISPR single guide RNA (MAST1-KI-sgRNA2), the single-stranded oligodeoxynucleotide (ssODN) donor used for homology-directed repair (HDR), and the forward and reverse primers used for genotyping PCR. **(B)** Schematic of the human

MAST1 locus (NM_014975.3) showing exons 23-26. The expanded sequence below indicates the sgRNA target site (red) and the SpCas9 protospacer-adjacent motif (PAM, NGG; green). **(C)** Representative Sanger sequencing chromatograms of the isogenic control line iCtrl (H9), the heterozygous clones Mut1 (Het21) and Mut2 (Het73), and a homozygous clone. The boxed nucleotide indicates the edited position (c.3889G>T, p.Asp1297Tyr): a wild-type G in iCtrl, overlapping G and T peaks in the heterozygous clones, and a single T peak in the homozygous clone.

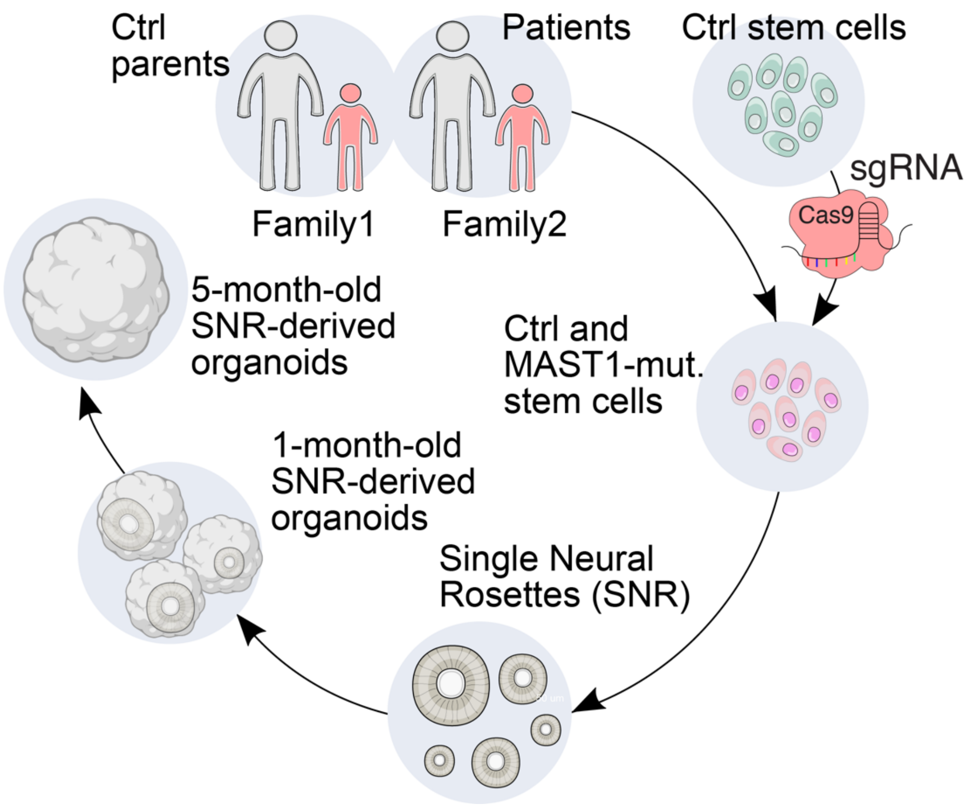


**Supplementary Figure 5. Schematic of the generation of iPSC-derived and CRISPR/Cas9-engineered organoids.** iPSCs were derived from patients with MAST1 mutations and from their unaffected parents, who served as Control 1 and Control 2, from Family 1 and Family 2 respectively. In parallel, the patient MAST1 mutation was introduced into control stem cells (H9) using CRISPR/Cas9 to generate isogenic mutant lines. Control and MAST1-mutant stem cells were differentiated into single neural rosettes (SNRs), which were then grown into SNR-derived organoids and analyzed at 1 month and 5 months.

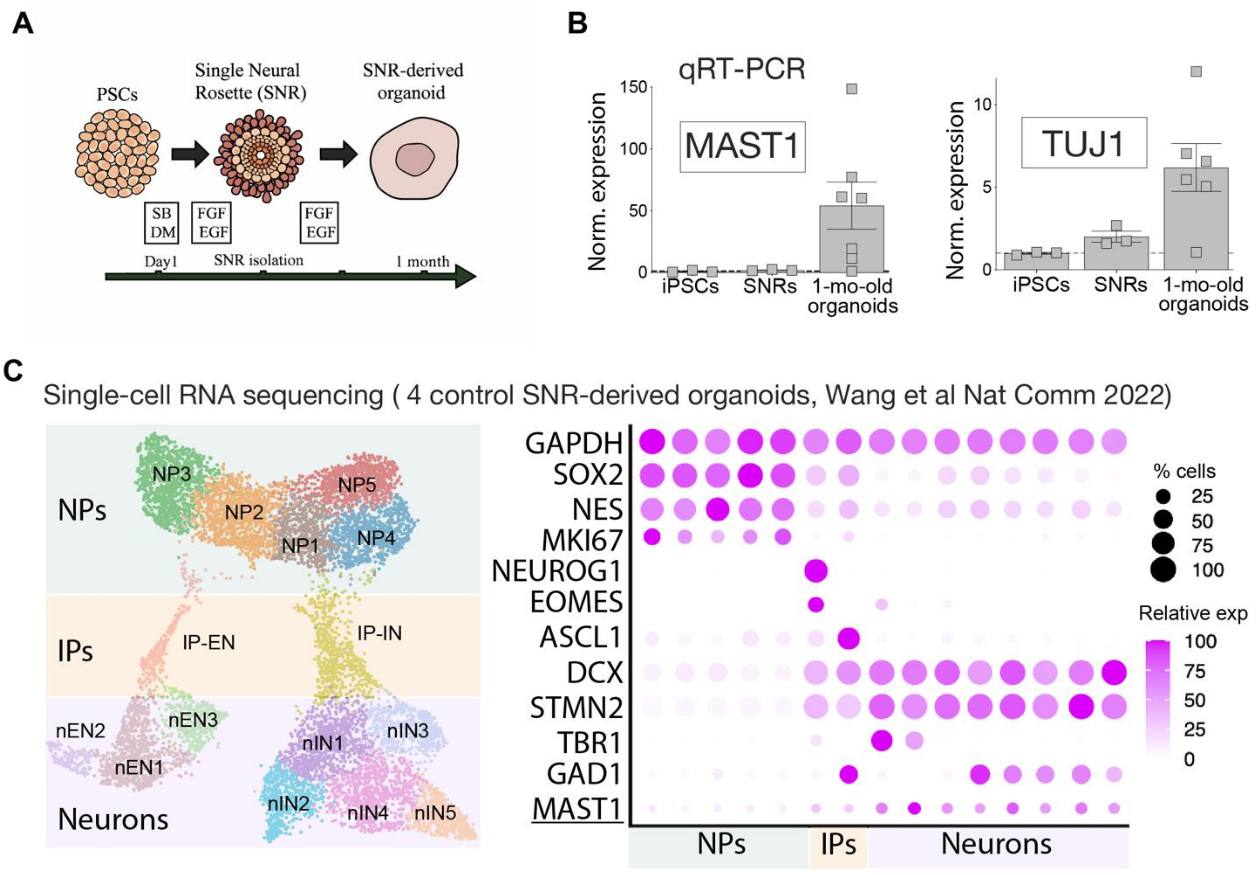


**Supplementary Figure 6. MAST1 expression in iPSCs, SNRs, and organoids. (A)** Schematic of SNR-derived organoid generation. Pluripotent stem cells (PSCs) were treated with SB431542 and dorsomorphin (SB/DM) from day 1, followed by FGF and EGF. SNRs were isolated and cultured in FGF/EGF to form organoids, which were analyzed at 1 month. **(B)** qRT-PCR analysis of *MAST1* and *TUJ1* mRNA expression in iPSCs, SNRs, and 1-month-old organoids. Expression was normalized to *GAPDH*. Both *MAST1* and *TUJ1* expression levels were increased in 1-month-old organoids. Each symbol represents one biological sample: n=3 independent wells of a 6-well plate for iPSCs, n=3 independent wells of a 6-well plate for SNRs, and n=6–7 organoids from 3 independent batches. Error bars = mean ± SEM. **(C)** Single-cell RNA-sequencing of four control SNR-derived organoids (data from Wang et al., 2022). Left, UMAP showing clusters of neural progenitors (NP1-NP5), intermediate progenitors of the excitatory (IP-EN) and inhibitory (IP-IN) lineages, and excitatory (nEN1-nEN3) and inhibitory (nIN1-nIN5) neurons. Right, dot plot of marker genes for progenitors (*SOX2*, *NES*, *MKI67*), intermediate progenitors (*NEUROG1*, *EOMES*, *ASCL1*), neurons (*DCX*, *STMN2*), excitatory neurons (*TBR1*) and inhibitory neurons (*GAD1*), with *GAPDH* as a reference. Dot size indicates the percentage of cells expressing each gene, and color indicates relative expression. *MAST1* expression is enriched in neuronal clusters.

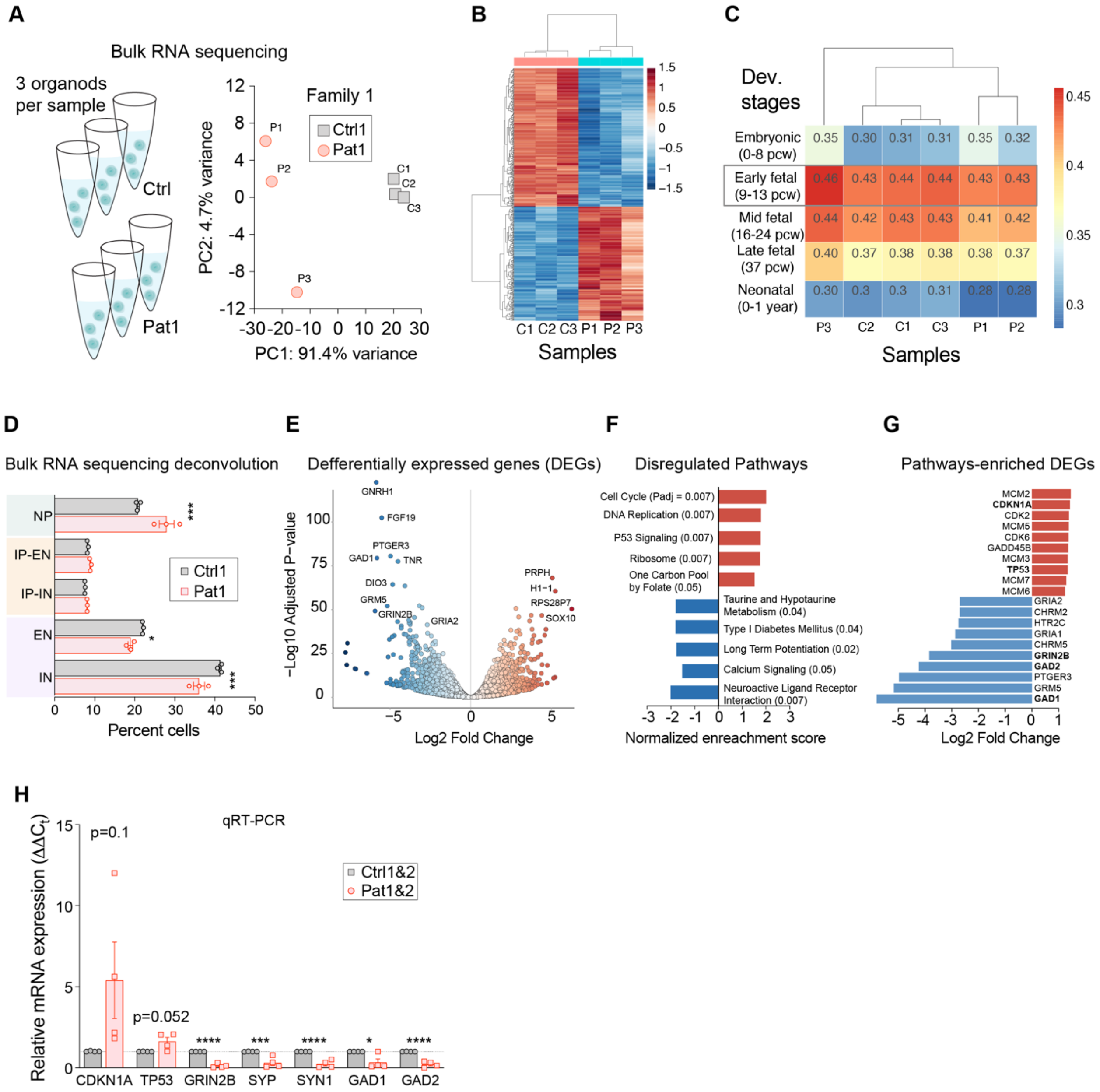


**Supplementary Figure 7. Transcriptomic profiling of patient-derived organoids and qRT-PCR validation. (A)** Left, bulk RNA-sequencing design: three organoids were pooled per sample, with three samples each from Ctrl1 (c1-c3) and Pat1 (p1-p3) of Family 1. Right, principal component analysis separating control and patient samples along PC1 (91.4% of variance; PC2, 4.7%). **(B)** Heatmap with clustering of differentially expressed genes (DEGs) between Ctrl1 and Pat1 organoids. The color scale indicates row-scaled expression (z-score). **(C)** Correlation of organoid transcriptomes with human brain developmental stages: embryonic (0-8 post-conception weeks, pcw), early fetal (9-13 pcw), mid fetal (16-24 pcw), late fetal (37 pcw), and neonatal (0-1 year). All samples correlated most strongly with the early fetal stage (boxed). **(D)** Cell type proportions calculated by deconvolution of the bulk RNA-sequencing data. Pat1 organoids contained a higher proportion of neural progenitors (NPs) and fewer excitatory neurons (ENs) and inhibitory neurons (INs) than Ctrl1. IP-EN and IP-IN, intermediate progenitors of the excitatory and inhibitory lineages. **(E)** Volcano plot of DEGs in Pat1 versus Ctrl1 organoids. Upregulated genes are shown in red and downregulated genes in blue; selected genes are labeled. **(F)** Gene set enrichment analysis of Kyoto Encyclopedia of Genes and Genomes (KEGG) pathways. Cell cycle, DNA replication, p53 signaling, ribosome, and one-carbon pool by folate were upregulated in Pat1 organoids, whereas neuroactive ligand–receptor interaction, long-term potentiation, calcium signaling, type I diabetes mellitus, and taurine and hypotaurine metabolism were downregulated. Adjusted P values are shown in parentheses. **(G)** Log2 fold change values for DEGs within the enriched pathways, including upregulated cell cycle and p53 pathway genes (*MCM2, MCM3, MCM5, MCM6, MCM7, CDK2, CDK6, CDKN1A, GADD45B, TP53*) and downregulated synaptic and neurotransmitter receptor genes (*GRIA1, GRIA2, GRIN2B, GRM5, CHRM2, CHRM5, HTR2C, PTGER3, GAD1, GAD2*). **(H)** qRT-PCR validation of selected DEGs in control (Ctrl1 and Ctrl2) and patient (Pat1 and Pat2) organoids. Relative mRNA expression was calculated using the $^{\Delta\Delta}$Ct method and normalized to *GAPDH* and to controls. Each symbol represents individual organoids (n=4). Error bars = mean ± SEM. *P<0.05, **P<0.01, ***P<0.001, and ****P<0.0001; unpaired two-tailed t-test.

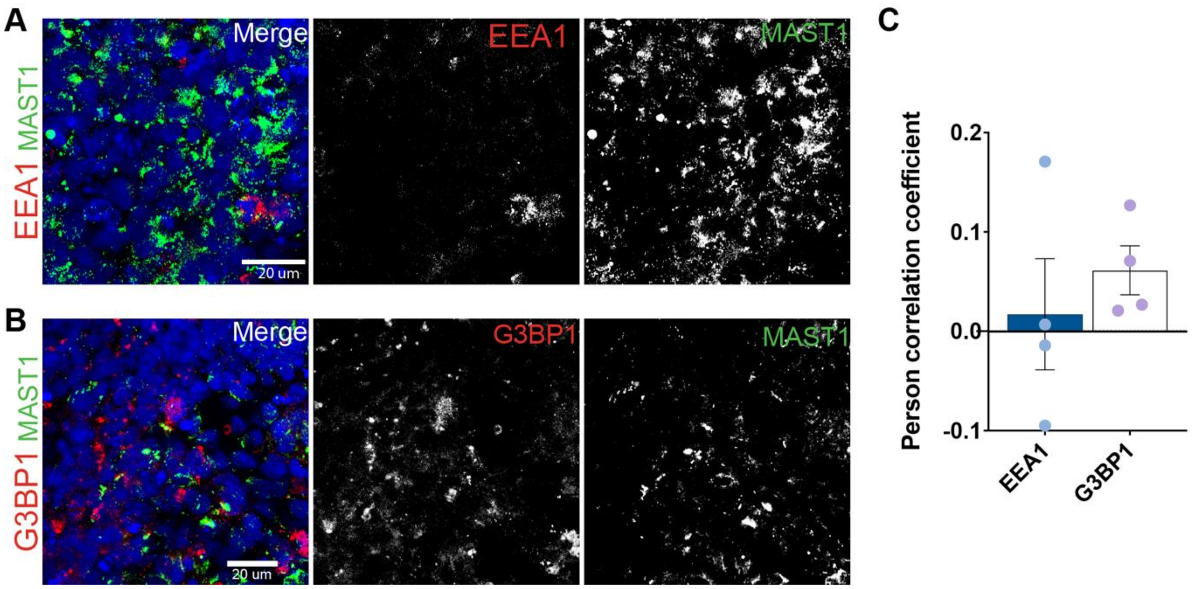


**Supplementary Figure 8. MAST1 does not colocalize with endosomes or stress granules in 5-month-old organoids. (A and B)** Representative immunofluorescence images of 5-month-old organoids stained for MAST1 (green) and early endosome marker EEA1 (red; A) or the stress granule marker G3BP1 (red; B). Nuclei are stained with DAPI (blue). Merged images (left) and single channels (grayscale) are shown. Scale bars, 20 µm. **(C)** Pearson correlation coefficients for MAST1 with EEA1 or G3BP1. Values near zero indicate the absence of colocalization. Each dot represents organoids (n=4). Error bars = mean ± SEM.

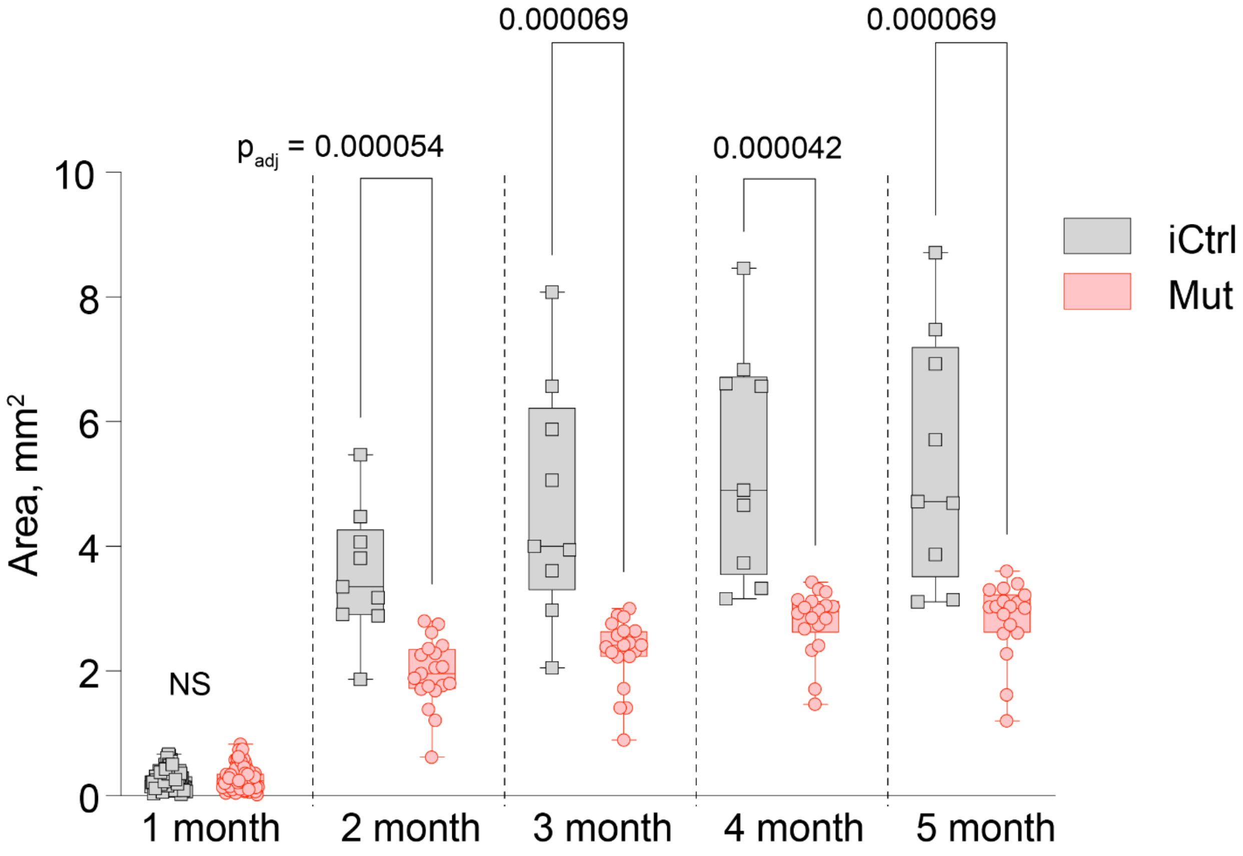


**Supplementary Figure 9. Growth of control and MAST1-mutant organoids over time.** Organoids size; area ($mm^2$) of isogenic control (iCtrl, gray) and MAST1-mutant (Mut, red) organoids measured monthly from 1–5 months. Organoid size was comparable at 1 month, but mutant organoids were significantly smaller than control organoids from 2 months onward. Each symbol represents one organoid. Adjusted P values are shown above each; Error bars = mean ± SEM. ns, not significant; unpaired two-tailed t-test.
